# Prophylactic TLR9 stimulation reduces brain metastasis through microglia activation

**DOI:** 10.1101/533927

**Authors:** Amit Benbenishty, Meital Gadrich, Azzurra Cottarelli, Alisa Lubart, David Kain, Malak Amer, Lee Shaashua, Ariella Glasner, Neta Erez, Dritan Agalliu, Lior Mayo, Shamgar Ben-Eliyahu, Pablo Blinder

## Abstract

Brain metastases are prevalent in various types of cancer, and are often terminal given low efficacy of available therapies. Therefore, preventing them is of outmost clinical relevance and prophylactic treatments are perhaps the most efficient strategy. Here, we show that systemic prophylactic administration of a TLR9 agonist, CpG-C, is effective against brain metastases. Acute and chronic systemic administration of CpG-C reduced tumor cell seeding and growth in the brain in three tumor models in mice, including metastasis of human and mouse lung cancer, and spontaneous melanoma-derived brain metastasis. Studying mechanisms underlying the therapeutic effects of CpG-C, we found that in the brain, unlike in the periphery, NK cells and monocytes are not involved in controlling metastasis. Next, we demonstrated that the systemically administered CpG-C is taken up by endothelial cells, astrocytes, and microglia, without affecting blood-brain barrier integrity and tumor brain extravasation. *In vitro* assays pointed to microglia, but not astrocytes, as mediators of CpG-C effects through increased tumor killing and phagocytosis, mediated by direct microglia-tumor contact. *In vivo*, CpG-C-activated microglia displayed elevated mRNA expression levels of apoptosis-inducing and phagocytosis-related genes. Intravital imaging showed that CpG-C-activated microglia cells contact, kill, and phagocytize tumor cells in the early stages of tumor brain invasion more than non-activated microglia. Blocking *in vivo* activation of microglia with minocycline, and depletion of microglia with a colony-stimulating factor 1 inhibitor, indicated that microglia mediate the anti-tumor effects of CpG-C. Overall, the results suggest prophylactic CpG-C treatment as a new intervention against brain metastasis, through an essential activation of microglia.

**Summary:** Brain metastases are prevalent and often terminal. Thus, reducing their occurrence could markedly improve cancer outcome. We show that systemic prophylactic and perioperative administration of a TLR9 agonist, CpG-C, reduced metastatic growth in experimental and spontaneous brain metastasis models, employing mouse and human tumors. CpG-C was taken up in the brain, without affecting blood-brain barrier integrity and tumor extravasation. *In vitro* assays, imaging flow cytometry, and intravital imaging pointed to microglia as mediators of CpG-C effects through contact-dependent tumor killing and phagocytosis; corresponding with *in vivo* mRNA profile. *In vivo* depletion studies proved that microglia, but not NK cells or monocytes, mediated the beneficial effects of CpG-C; Also hindered by blocking microglial activation. In-toto, perioperative treatment with CpG-C should be considered clinically relevant.

**Significance:** Preventing brain metastases is paramount, as they are considered incurable and their incidence is on the rise due to prolonged survival of cancer patients. Here, we demonstrate that systemic prophylactic treatment with CpG-C reduces peripheral and brain metastasis of mouse and human lung cancers. While traditional therapies are halted during the perioperative period, we found systemic CpG-C treatment during this time frame beneficial in a model of spontaneous brain metastases following excision of a primary melanoma tumor, comprehensively mimicking the clinical setting. Mechanistically, we show microglia activation with CpG-C results in tumor cell eradication, pointing to microglia as potential therapeutic targets. Importantly, CpG-ODNs have negligible toxicity in humans. Therefore, CpG-C may be used prophylactically and during the perioperative period in high-risk cancers.

## Introduction

Ten to twenty percent of cancer patients develop brain metastases, commonly as the final stage of cancer progression, with lung and melanoma cancers having the highest incidence (40-50% and 30-50%, respectively) (1). Therapies include surgery and radiation, however both treatments result in only a modest survival advantage and are associated with cognitive impairments (2). Chemotherapy is often inefficient due to impermeability of the blood-brain barrier (BBB) (1), and as it often induces astrocyte-derived tumor-protecting responses (3). Overall, the efficacy of currently available treatments for brain metastasis is extremely limited, making it a deadly disease with a short survival period (2). Thus, prophylactic approaches against the establishment of brain metastasis, or early elimination of brain micrometastases, could prove key in treating cancer (2,4,5), even more so given ongoing progression in early cancer detection and prevention of peripheral metastases.

In recent years, immune-modulation using toll-like receptors (TLRs) agonists has been given much attention as a therapeutic approach against primary tumors and metastasis (6). Specifically, the TLR9 agonists CpG-oligodeoxynucleotides (ODNs) are being explored in a wide range of tumor types, both as single agents and as adjuvants (7,8), and are being tested in several clinical trials (9). In various animal models, CpG-ODN treatment was shown to reduce mammary lung metastases by eliciting anti-tumor NK activity (9), and even result in rapid debulking of large tumors by macrophage stimulation (10). Employed prophylactically, CpG-ODNs were shown to markedly improve resistance to experimental and spontaneous peripheral metastasis of mammary (11), colon (12), and melanoma (13) tumors.

Given the low success rate of treatments against established brain metastases (1), prophylactic treatment against metastatic brain disease may be key to improve survival rates (4). Such treatment should be given chronically between primary tumor diagnosis and until several days/weeks following tumor removal. This time frame includes the short perioperative period, which was shown to constitute a high-risk period for initiation or accelerated progression of metastasis (14). Prophylactic treatment should be especially advantageous in patients with primary tumors that have high potential of developing brain metastases, such as lung, melanoma and breast cancers (15). In fact, the concept of prophylactic treatment against brain metastasis is not unprecedented and is routinely practiced in the clinic. Small-cell lung cancer (SCLC) patients without detectable brain metastases often undergo prophylactic whole brain radiation therapy, thereby reducing occurrence of brain metastases and improving survival (16,17). However, to implement a prophylactic approach against brain metastases in a wider range of patients, a less toxic (18) treatment is required. TLR9 stimulation using CpG-ODNs is particularly well suited to meet this need as, it has negligible toxicity in humans (19–21), and has already promising preclinical outcomes in other organs (10–13); therefore, it should also be considered a potential prophylactic approach against the establishment of brain metastases.

In the brain, TLR9 is expressed on neurons, astrocytes, microglia, and endothelial cells (22,23). Recent studies suggest that TLR9 signaling plays a key role in cerebral ischemia (24), cerebral malaria (25), Alzheimer’s (26,27), and seizures (28), pointing to its key role in healthy brain function and neuro-immune modulation. Notably, intracerebral (29,30) and retro-orbital (31) administration of CpG-ODNs were shown to hinder growth of glioma (31) and intracranially-injected melanoma cells (29,30). Importantly, CpG-ODN yielded promising initial outcomes with minimal toxicity in a few phase I/II clinical trials of recurrent (20,21) and de novo (19) glioblastoma, when injected into tumor-excised lesions. However, as a prophylactic measure against potential brain metastasis, CpG-ODNs would need to be administered systemically, provided they can cross the BBB.

Here, we assessed the efficacy of a systemic administration of CpG-C as a prophylactic treatment for brain metastasis using three pre-clinical mouse models, including experimental metastasis of syngeneic and of human lung carcinoma, and spontaneous metastasis of syngeneic melanoma. Importantly, the inoculation methodologies and imaging approaches implemented here preserve intact both the neuro-immune niche and brain hemodynamics intact; crucial factors affecting metastatic early stages (32). We demonstrate that acute and chronic prophylactic treatments result in reduced brain metastasis in both sexes and across ages. Notably, we found that NK cells and monocytes/macrophages do not take part in the initial steps of the metastatic process in the brain, nor do they mediate the effects of CpG-C, as opposed to their role in peripheral organs. We establish that peripherally administered CpG-C crosses into the brain parenchyma without affecting BBB permeability, and that cerebral endothelial cells, astrocytes, and microglia uptake it. We found that CpG-C-activated primary microglial cells and the N9 microglial cell line (but not primary astrocytes) eradicate tumor cells *in vitro*, through direct contact, by increasing microglia cytotoxicity and phagocytic potential. Importantly, we demonstrate *in vivo* that following systemic CpG-C treatment microglia cells contact, kill, and phagocytize tumor cells during the early stages of invasion into the brain. Blocking microglia activation or depleting them abolished the beneficial effects of CpG-C. Taken together, our results point to CpG-C as an important potential prophylactic treatment against brain metastasis through direct activation of microglia.

## Methods

### Cell preparation and *in vitro* experiments

<u>Tumor cells</u> – Mouse D122 Lewis lung carcinoma (LLC) and mouse Ret melanoma cells(33) (both syngeneic to C57BL/6J mice), and human PC14-PE6 adenocarcinoma cells were cultured in complete media (RPMI1640 supplemented with 10% fetal bovine serum and 1% penicillin/streptomycin; Biological Industries). D122-LLC and PC14-PE6 cells (kindly provided by Prof. Isaiah Fidler) were double labeled with mCherry and Luc2 (pLNT/Sffv-MCS/ccdB plasmid was kindly provided by Prof. Vaskar Saha), and Ret melanoma cells were labeled with mCherry. For two-photon experiments, D122 cells were infected with pLVX-tdTomato-N1 (Clontech). For experiments assessing cancer cell retention, cultured tumor cells were incubated with ^125^IUDR during the last 24 hours of proliferation. Before injection, cells were washed and harvested (0.25% trypsin-EDTA; Invitrogen) at ~90% confluence, re-suspended in PBS supplemented with 0.1% BSA (Biological Industries), and kept on ice throughout the injection procedures, completed within 3 hours of cell harvesting. More than 95% of cells were vital throughout the injection period.

<u>Primary cultures</u> (Fig. 5a-d) – We followed the mild trypsinization procedure (34). Briefly, cortices of 1-3 days old C57BL/6J pups were cultured in 12-well plates at a concentration of 4▯10^5^ cells/well. After 18-25 days, astrocytes were removed with trypsin and cultured on separate plates. Cultures were used within 4 days from trypsinization.

<u>Microglial N9 cell line</u> (35) (Fig. 5e-i) – cells were cultured in complete media (see above) at a concentration of 4▯10^4^ cells/well.

#### Experimental procedures

Cultures were subjected to 100nM/L of CpG-C or non-CpG ODN (control) for 24 hours and media was harvested for conditioned-media experiments (Fig. 5b,d,g). Cultures were washed twice and supplied with fresh media. For contact co-cultures experiments (Fig. 5a,c,e,h), D122 cells were plated on top of the microglia cultures for six hours, following which cell-lysis (primary cultures; Fig. 5a,c), bioluminescence imaging (N9 cultures; Fig. 5e), or FACS analysis (Fig. 5h) assays were conducted. For no-contact co-cultures (Fig. 5i), D122 cells were plated on 13mm cover slips, and placed on top of 2mm thick costume made polydimethylsiloxane 11mm rings over the microglial cultures (sharing the same media for six hours). For conditioned-media experiments (Fig. 5f), D122 cultures were washed and supplied with fresh or conditioned-media harvested from microglia or astrocyte cultures, for six hours.

##### Cell-lysis assay

(Fig. 5a-d) – Standard cytotoxicity assay was conducted as previously described (36). Briefly, we used two concentrations of ^125^IUDR-lebeled D122 cells in 12-well plates (1×10^4^ and 2×10^4^ cells/well). Radioactive signal from the media was measured using a gamma counter (2470, PearkinElmer). Specific killing was calculated as:

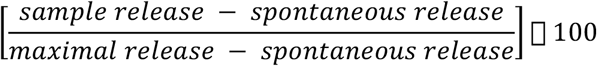

##### *In vitro* bioluminescence viability assay

(Fig. 5e-g) – N9 cells were plated in 24-well plates (40×10^3^ cells/well) and treated with CpG-C or non-CpG ODN for 24 hours. We used two concentrations of Luc2-labeled D122 cells in 24-well plates (1.6×10^4^ and 3.2×10^4^ cells/well). D-luciferin (30mg/ml, 10μl) were mixed in each well and bioluminescence signal was immediately measured for two minutes using Photon Imager and analyzed with M3 Vision (Biospace Lab).

Lysis and bioluminescence assessments were repeated in at least three separate experiments, each one conducted in quadruplicates or more.

##### Apoptosis quantification

(Fig. 5h and Supplementary Fig. 8) – co-cultures of N9 and D122 cells were stained for annexin V (88-8005-72, eBiosciece), as per manufacturer’s instructions. We quantified the percent of annexin V positive (apoptotic) cells from all mCherry positive (D122) cells using FACScan (Becton Dickinson).

##### Phagocytosis assay

(Fig. 5i) – N9 cells were plated in 96-well plates (30×10^3^ cells/well) and treated with CpG-C or non-CpG ODN for 24 hours. Cultures were washed twice and plated with pHrodo™ Red Zymosan Bioparticles™ (ThermoFisher Scientific) conjugate for phagocytosis, according to the manufacturer’s instructions. These particles become fluorescent only after phagocytized into the lysosomes. Fluorescence was measured with Synergy HT (BioTek) microplate reader at 545/585 (Ex/Em) every 30-60 minutes thereafter (up to 6 hours). The maximum difference between experimental groups was used for statistical analysis.

##### Scratch assay

(Supplementary Fig. 7a) - N9 cells were plated in 96-well plates (30×10^3^ cells/well) and treated with CpG-C or non-CpG ODN as above. Plates were washed and fresh media was added. Confluent cultures were scratched (700μm) using the IncuCyte Zoom system (Essen BioScience), washed, and imaged once every two hours for 28 hours.

### Animals and Anesthesia

All studies were approved by the Tel Aviv University and Columbia University corresponding ethics committees for animal use and welfare and in full compliance with IACUC guidelines. C57BL/6J, athymic nude mice (Hsd:Athymic Nude-Foxn1nu), CX3CR1^GFP/+^ knock-in (B6.129P-Cx3cr1tm1Litt/J), and Tg eGFP-Claudin5 (37) male and female mice were used (8-52 weeks old; age matched within experiment). Animals were housed under standard vivarium conditions (22±1 °C, 12h light/dark cycle, with ad libitum food and water). Anesthesia was first induced by 5% Isoflurane, and then maintained on 1.5-2% throughout the procedures. When anesthetized, core body temperature of animals was maintained at 37°C.

### Internal carotid artery inoculation of tumor cells

Tumor cells were injected using the assisted external carotid artery inoculation (aECAi; Fig. 1a), as previously described (32). Briefly, mice were anesthetized and the external carotid artery (ECA) uncovered. A 6-0 silk-suture ligature was loosely placed around the ECA proximal to the bifurcation from the common carotid artery (between the superior thyroid artery and the bifurcation). A second ligature was tied on the ECA distal to the bifurcation. A NANOFIL-100 (WPI) syringe with a 34G beveled needle was mounted to a micromanipulator (M33, Sutter Inc). The needle was inserted slowly into the lumen of the ECA and advanced to the point of bifurcation. The first ligature was tied around the needle, and 1×10^5^ cells in PBS (100μl) were slowly infused into the internal carotid artery. The needle was then removed, the ligature quickly tied, and the skin sutured. Total time for the complete procedure is ten minutes.

**Figure 1:**
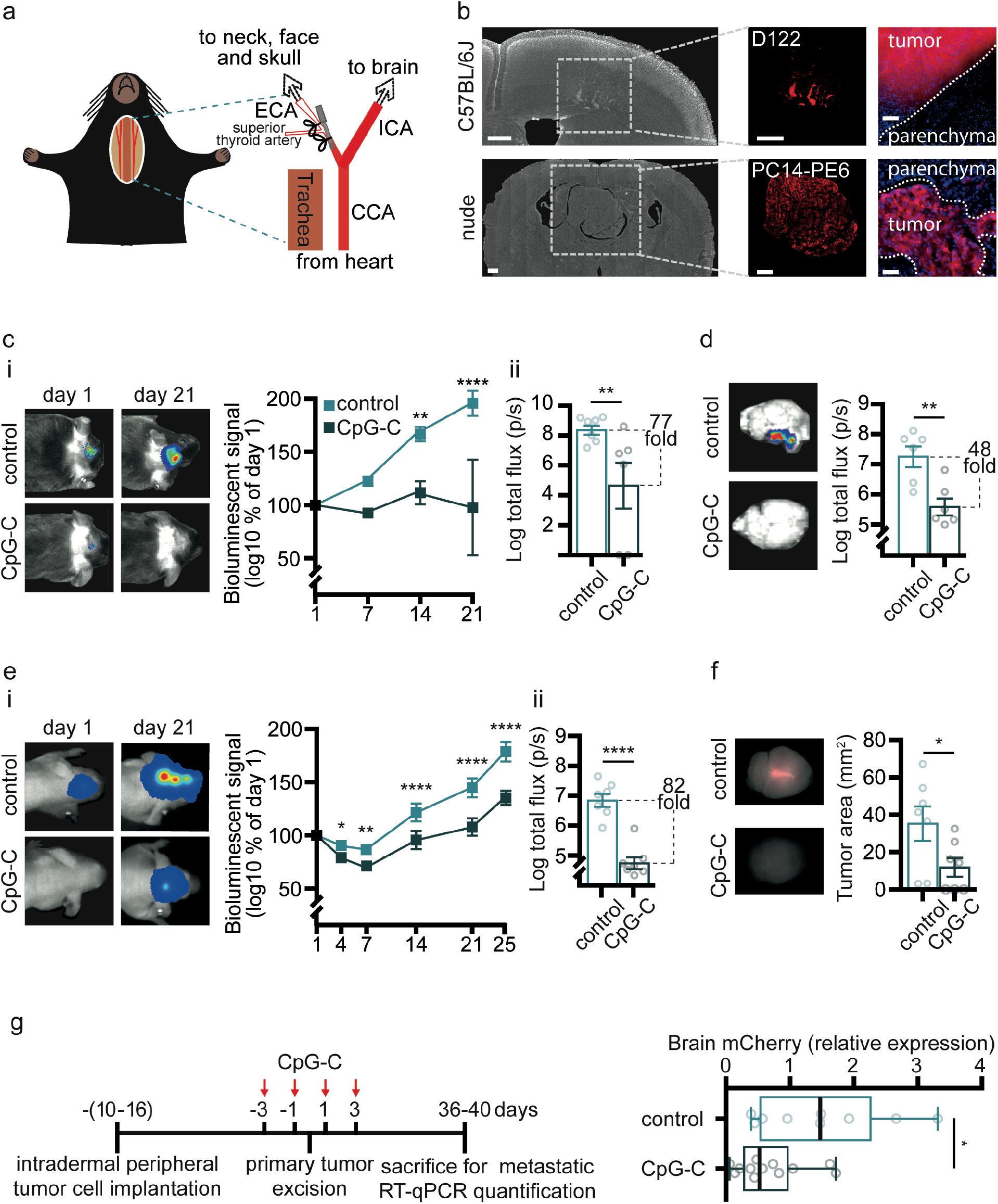
A single systemic prophylactic treatment with CpG-C results in long term reduction of metastatic burden in the brain. (**a**) In the experimental metastasis models, we used the assisted external carotid artery inoculation (aECAi) approach (32) for injection of tumor cells (see *Methods*); a method that improves brain targeting and preserves cerebral hemodynamics. (**b**) Histological images of D122 brain metastases from C57BL/6J mice on day 21 post tumor inoculation show well-demarcated metastases, as well as vessel co-option growth. PC14-PE6 brain metastases from nude animals on day 25 post tumor inoculation show large well-demarcated metastases. Scale bar is 500μm for the images on the left and middle, and 50μm for the images on the right. (**c-f**) A single prophylactic systemic (i.p.) injection of 4mg/kg CpG-C resulted in reduced growth of experimental brain metastases, as indicated by bioluminescence and fluorescence imaging. (c) C57BL/J6 mice injected with syngeneic D122 tumor cells, and pre-treated with CpG-C, had reduced tumor burden compared to control animals, becoming significant on day 14 (**ci**; n=6-7; F_(1,11)_=19.02, p=0.0011), and reaching a 77-fold difference in total flux on day 21 (two-tailed Mann-Whitney U=3, p=0.0082; **cii**). Interestingly, in two CpG-C-treated animals bioluminescent signal gradually decreased and disappeared on day 21. (**d**) *Ex-vivo* bioluminescence imaging of the brains from the syngeneic model indicated a 48-fold reduced tumor burden (total flux) in CpG-C treated animals (n=6; two-tailed unpaired student t-test, t(10)=3.722, p=0.0040). (**e**) Athymic nude mice injected with human (xenograft) PC14-PE6 tumor cells, and pre-treated with CpG-C, had reduced tumor burden compared to control animals, becoming significant on day 4 (**e.i**; n=7; F_(1,12)_=77.45, p<0.0001), and reaching a 82-fold difference in total flux on day 25 (two-tailed unpaired student t-test, t(12)=7.09, p<0.0001; **e.ii**). (**f**) Using Maestro fluorescence imaging, a reduction in brain tumor burden (i.e. tumor area) was evident in the human xenograft model in CpG-C-treated animals (n=7; two-tailed Mann-Whitney U=8, p=0.0373). (**g**) (**left**) timeline for spontaneous melanoma brain metastasis model (33) (see *Methods*). (**right**) CpG-C treatment during seven perioperative days resulted in reduced micrometastases in the brain (measured by mCherry mRNA expression; n=9 and n=12 for control and CpG-C, respectively; two-tailed unpaired student t-test, t(19)=2.278, p=0.0345). Background of images was manually removed. Boxplot whiskers represent min-max range.

### Spontaneous melanoma brain metastasis

For a spontaneous brain metastasis model (Fig. 1g) we used the Ret-melanoma model, we have recently established and validated (33). Briefly, mice were anesthetized by isoflurane, and a total of 5×10^5^ (50μl) Ret-mCherry sorted (RMS) cells in a 1:1 suspension of PBS with growth factor–reduced Matrigel (356231, BD Biosciences) were injected subdermally to the right dorsal side, rostral to the flank, with a 29G insulin syringe (BD Biosciences). Tumors were measured four times weekly by calipers. Tumor volumes were calculated using the formula X^2^×Y×0.5 (X-smaller diameter, Y-larger diameter). We aimed to remove the tumor at a size of ~500 mm^3^. Therefore, and based on our experience, once tumors reached a size of ~125mm^3^, mice were given two injections of CpG-C or PBS (control group) every other day. One day later (i.e. three days following the first CpG-C treatment and one day following the second CpG-C treatment), tumor sizes were verified (meeting our expectations, with no differences between treatment groups), and immediately removed. The last tumor removal was carried out six days after the first removal. For tumor excision, mice were anesthetized with isoflurane, and an incision, medial to the tumor, was made in the skin. Tumors were detached from inner skin with clean margins to prevent recurrence. Tumor-associated connective tissue and blood vessels were detached, and the incision was sutured. Primary tumors were sectioned and measured with no difference identified at excision time (Supplementary Fig. 1a). Mice were weighed weekly and monitored for relapse. Nine weeks of last tumor excision, mice were deeply anesthetized with isoflurane and transcardially perfused with cold PBS. Brains and lungs were harvested, macroscopically examined for abnormal lesions and flash-frozen in liquid nitrogen. RNA was isolated using EZ-RNA II kit (20-410-100, BI) according to the manufacturer’s instructions. Whole organs were homogenized in denaturation solution A in M tubes (130-096-335, Milteny Biotec) by gentleMACS dissociator (Milteny Biotec). Reverse transcription was performed with qScript (95047-100, Quanta Biosciences). qRT-PCR analyses were conducted using PerfeCTa SYBR Green FastMix, ROX (95073-012- 4, Quanta Biosciences) with primers for Hprt (F sequence – GCGATGATGAACCAGGTTATGA; R sequence – ATCTCGAGCAAGTCTTTCAGTCCT) and mCherry (F sequence – GAACGGCCACGAGTTCGAGA; R sequence – CTTGGAGCCGTACATGAACTGAGG). In all analyses expression results were normalized to *Hprt*. RQ (2^-ΔΔCt^) was calculated.

Of the 50 animals initially injected with tumor cells, two animals did not develop primary tumors and were withdrawn from the experiment; of the remaining 48 animals, 28 animals were treated with CpG-C and 20 with PBS (control). Twenty-two animals (45% of control and 46% CpG-C treated) died during the period between tumor excision and the day of sacrifice, leaving 15 CpG-C treated animals and 11 control animals. In three CpG-C animals and two control animals we did not detect mCherry RNA in the brains. The herein development of primary tumor and metastases is expected based on our previous studies in this tumor model (33). Tumor burden was compared in animals bearing brain micrometastasis.

#### Oligodeoxynucleotides (ODN) treatment

CpG-C, CpG-C-FITC, and CpG-C-TAMRA (ODN 2395: 5′-TCGTCGTTTTCGGCGCGCGCCG-3′) with a phosphorothioate backbone and non-CpG ODN (ODN 2137: 5′-TGCTGCTTTTGTGCTTTTGTGCTT- 3′), endotoxin free, were purchased from Sigma-Aldrich. Two different controls were used: Phosphate buffered saline (PBS), and non-CpG ODN, which lacks C-G motifs (counterbalanced within experiments with no differences in results). Both CpG-C variants and non-CpG ODN were diluted in PBS, and administered intraperitoneally (100μl) at a dose of 0.4 or 1.2mg/kg (Supplementary Fig. 2c), or 4mg/kg (all *in vivo* experiments). No differences were found between PBS and non-CpG ODN treated animals and therefor combined in the statistical analyses (Supplementary Fig. 6a).

#### Depletion of NK cells and monocytes/macrophages

For depletion of NK cells (Fig. 2a,b), anti-NK1.1 monoclonal antibodies (mAbs) were intraperitoneally administered (4mg/kg) twenty-four hours before tumor cell injection. 12E7 mAb against human CD99 served as control. Antibodies (38) were kindly provided by Prof. Ofer Mandelboim (The Hebrew University of Jerusalem, Israel). To verify depletion of NK cells, blood was collected from animals when sacrificed, and prepared for staining with NK1.1 FITC (eBioscience) and NKp46 PE (BioGems) (39). FACS analysis indicated >90% depletion (Supplementary Fig. 3a).

**Figure 2:**
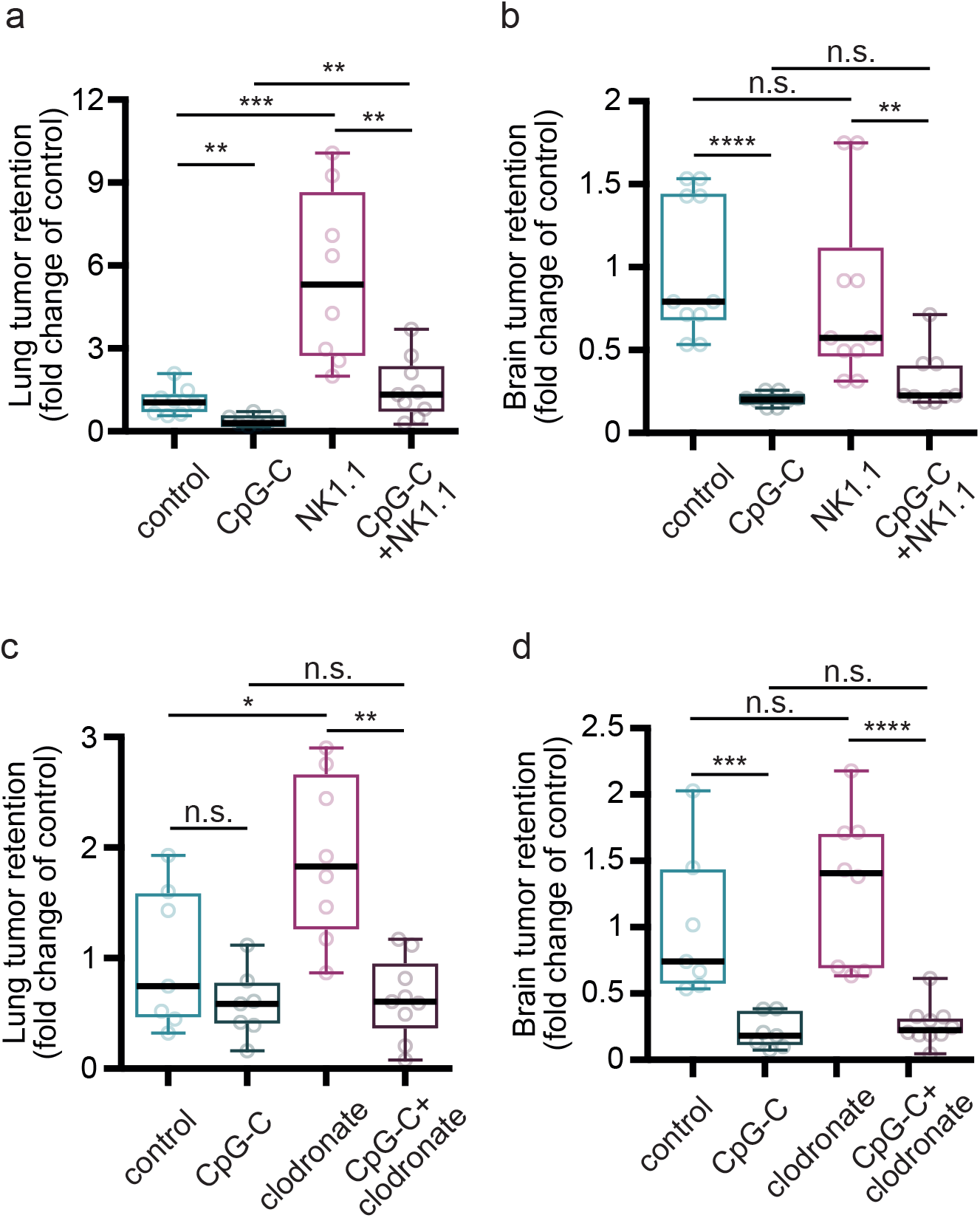
The effects of CpG-C on brain metastases are not mediated by NK cells or monocytes. (**a**) Depletion of NK cells using NK1.1 antibody, resulted in a 5-fold elevation in D122 Lewis lung carcinoma tumor retention in the lungs (n=8; t(14)=4.4781, p=0.0001), and partially blocked the beneficial effects of CpG-C (n=8; t(14)= 1.1517, p=0.0038), evident in naïve animals (n=8; t(14)=0.7002, p=0.0019). (**b**) In brains of the same animals, NK depletion had no effect on tumor retention (n=8; t(14)=0.1894, p=0.3935), nor mediated the beneficial effects of CpG-C (n=8; t(14)=0.1099, p=0.0811), evident in both naïve (n=8; t(14)=0.7973, p<0.0001) and NK depleted animals (n=8; t(14)=0.4979, p=0.0056). (**c**) Depletion of monocytes using clodronate liposomes resulted in increased lung tumor retention of D122 Lewis lung carcinoma cells (n=7-8; t(13)=0.9072, p=0.0292), an effect rescued by CpG-C (n=8-9; t(15)=1.270, p=0.0003), indicating lung tumor retention is mediated by monocytes, while they do not mediate the effects of CpG-C. (**d**) In brains of the same animals, monocyte depletion did not affect tumor retention (n=7-8; t(13)=0.3028, p=0.3081), and CpG-C reduced tumor retention in naïve (n=7; t(12)=0.7910, p=0.0006), and in monocyte-depleted animals (n=8-10; t(16)=1.0377, p=0.0001). Two-way permutations were used for the above analyses. Boxplot whiskers represent min-max range.

For monocytes/macrophages depletion (Fig. 2c,d), we administered clodronate liposomes (ClodronateLiposomes.org) intravenously (200μl) twenty-four hours before tumor cell injection. PBS liposomes served as controls. To verify depletion of monocytes, but not microglia, blood and brains were collected from animals when sacrificed, and prepared for staining with F4/80 FITC and CD11b APC (BioGems). FACS analysis indicated >85% depletion of monocytes/macrophages, without affecting microglia viability (Supplementary Fig. 3b).

#### Microglia inactivation

(Fig. 7a,b) – To block microglia activation and transition into an inflammatory state (40), minocycline hydrochloride (Sigma-Aldrich) was administered intraperitoneally at a dose of 40mg/kg (200μl) at 48, 32, and 24 hours before tumor cell injection.

#### Depletion of microglia cells

(Fig. 7c) – For depletion of microglia cells mice were administered the dietary inhibitor of colony stimulating factor-1 receptor (CSF1R), PLX5622 (1200mg/kg chow; provided by Plexxikon Inc. and formulated in AIN-76A standard chow by Research Diets Inc.), for 18 days; resulting in near complete elimination of microglia cells (41). AIN-76A standard chow served as control (Research Diets Inc.).

### Histology

C57BL/6J and athymic nude mice from the bioluminescence experiments were perfused with PBS supplemented with 30U heparin (Sigma) and 4% PFA (EMS). Brains were harvested, fixed overnight in 4% PFA, and placed in 30% sucrose overnight. Thirty-micron sections (Leica SM 2000 microtome) were counterstained with DAPI (MP Biomedicals). Images of the sections were obtained using a fluorescent microscope (Olympus ix81; Fig. 1b).

To visualize CpG-C uptake in the parenchyma, TAMRA-labeled CpG-C was injected to CX3CR1^GFP/+^ mice. Twenty-four hours later animals were perfused, and brains fixed and sectioned. Astrocytes were stained using a primary anti-GFAP antibody (1:800; Invitrogen), and endothelial cells with anti-CD31 (PCAM-1) antibody (1:500; Santa Cruz). A secondary Alexa 647 antibody (1:600; Invitrogen) was used, coupled with DAPI (1:1000; ENCO) staining. Images of the sections were obtained using a Leica SP8 confocal microscope at 0.5μm intervals using a ×63 (NA – 1.4) oil immersion objective.

### Lysotracker staining

Cultures of N9 cells grown on cover slips were treated with CpG-C-TAMRA for 24 hours and washed three times. LysoTracker™ Blue DND-22 (50nM, ThermoFisher Scientific) was applied for 30 minutes at 37°C, and cover slips were washed and mounted on slides. For staining of CpG-C uptake *in vivo* a single cell suspension was prepared from CX3CR1^GFP/+^ mice treated with CpG-C-TAMRA as described in the ImageStream FACS analysis protocol herein. LysoTracker™ Blue DND-22 (50nM, ThermoFisher Scientific) was then mixed into the cell suspension for 30 minutes at 37°C and cells were mounted on a cover glass and imaged with a Leica SP8 confocal microscope using a ×63 (NA – 1.4) oil immersion objective (Fig. 5b). Similarly, for the cells extracted from the brains of animals, we imaged only GFP positive cells (i.e. microglia).

### Claudin5 continuity and IgG and biocytin-TMR leakage quantification

Tg eGFP-Claudin5 (37) mice were treated with CpG-C or PBS, and 23h later were injected with 1% biocytin-TMR (i.v., Life Technologies). One hour later animals were sacrificed and perfused with PBS and 4% paraformaldehyde (PFA). Brains and livers were harvested, fixed for six hours in 4% PFA at 4°C, and placed in 30% sucrose overnight at 4°C. Tissues were embedded in O.C.T (Sakura), sectioned (12μm) using a Leica cryostat, and stained for eGFP (1:1000; Life Technologies) and IgG (1:1000; Invitrogen). Z-stacks of the sections were obtained with a Zeiss LSM700 confocal microscope using a water immersion ×40 objective (NA – 1.2) and maximum projections were created using Fiji (version 1.0). At least 5 images were used for quantification for each anatomical region. Biocytin-TMR and IgG intensity was quantified using Fiji software and normalized on fluorescence intensity in the liver (Fig. 4a,b and Supplementary Fig. 4a-c). For quantification of gaps in tight junctions (Fig. 4c and Supplementary Fig. 4d), we quantified the percentage of junctional strands showing at least one gap (defined as a discontinuity in eGFP-Caudin5 signal>0.4μm) over the total number of junctional strands (37).

### Immune infiltration analysis

To test whether CpG-C affects immune cell infiltration into the brain, sections of PBS and CpG-C treated mice were stained for CD68 (1:1,000; Abcam) and CD4 (1:200; Abcam). To assure we do not analyze immune cells arrested in the vessels, we co-stained slices with laminin (1:1,000; Sigma) to detect vessel walls. As a positive control we used spinal cord sections of experimental autoimmune encephalomyelitis (EAE) mice (refer to (42) for experimental procedure; Fig. 4d).

#### *In vivo* and *ex-vivo* bioluminescent imaging

(Fig. 1c-e) – To follow progression of tumor growth *in vivo*, we used an IVIS SpectrumCT (PerkinElmer) for the syngeneic model and Photon Imager (Biospace Lab) for the xenograft model. Briefly, mice were anesthetized and injected with D122-mCherry-Luc2 (C57BL/6J) or PC14-PE6-mCherry-Luc2 (athymic nude) cells. Imaging sessions were conducted on days 1, 4, 7, 14 and 21 following tumor cell administration (in the xenograft model, also on day 25). After the last *in vivo* imaging session in the syngeneic model, mice were sacrificed, and brain and extra-cranial head tissue were rapidly imaged separately. Notably, tissue from one control animal was lost in the final process. Each imaging session was preformed between 10-20 minutes following D-Luciferin sodium salt injection (30mg/ml, 100μl, i.p; Regis Technologies), as this time frame exhibited maximal and steady intensity. Analysis was done using Living Image software (version 4.3.1) for the IVIS images data, and M3 vision for the Photon Imager data.

#### *Ex-vivo* fluorescence imaging

(Fig. 1f) – To quantify fluorescence in brains of athymic nude mice injected with PC14-PE6-mCherry-Luc2, animals were decapitated and brains were extracted immediately following the last imaging session. We used a Maestro spectral fluorescence imaging system (Cambridge Research & Instrumentation) and quantified fluorescent signal using Maestro version 2.2 software. Regions of interest (ROIs) were drawn on each fluorescent signal to quantify the area of fluorescent signal.

### Assessment of brain and peripheral organ retention of cancer cells

Mice were injected with 1×10^5 125^IUDR labeled D122-LLC cells using the aECAi approach (32), and euthanized 24 hours later. Animals were transcardially perfused with 20ml PBS supplemented with 30U heparin (Sigma-Aldrich). Brain and lungs were collected, and radioactivity was measured using a gamma counter (2470, PearkinElmer; Figs. 1f-h, 2, 7a,c).

### Two-photon laser scanning microscopy

For two-photon microscopy measurements, CX3CR1^GFP/+^ and WT mice were implanted with a polished and reinforced thin-skull (PoRTS) window, as previously described (43). Importantly, this craniotomy does not elicit an inflammatory response (43). Mice were then habituated to the imaging apparatus for 7 days to reduce procedural stress. CX3CR1^GFP/+^ animals were injected with 1×10^5^ tdTomato-labeled D122 cells. Before imaging, mice were injected with Alexa Fluor 633 hydrazide (2.5% w/v, i.v.; Invitrogen) for visualization of arteries (40). Imaging sessions were initiated 2-4 hours after tumor cell inoculation, and at days 1, 2, 4 and 7, returning to the exact same location each session. Imaging was conducted at depths of 50-200μm with a custom-modified two-photon laser-scanning microscope based on a Sutter MOM (Sutter Inc) controlled through the ScanImage software (Vidrio Technologies). A Chameleon Ultra II (Coherent Inc) provided the 80MHz, 140fs pulsed light used for imaging and laser photodamage.

For quantification of microglia-tumor cells interaction, 150μm stacks were obtained and max projected every 10μm. The number of contacts and internalization events in each stack were manually quantified blindly at 4 hours following tumor cell injection, and at days 1, 4 and 7 (Fig. 6d,e). For imaging CpG-C uptake by microglia *in vivo* (Supplementary Fig. 5a), baseline imaging of cortices of CX3CR1^GFP/+^ mice was performed at 890nm, CpG-C-TAMRA was injected (4mg/kg; 100μl; i.p), and 24h later mice were imaged again at the exact same locations (Supplementary Fig. 5a).

For longitudinal BBB assessment (Fig. 4e,f), WT mice implanted with a PoRTS window were treated with four PBS or CpG-C injections every other day (similarly to the spontaneous melanoma brain metastasis experiment). BBB leakage dynamics of a low molecular weight dye (sodium fluorescein; NaF; 376Da; Sigma-Aldrich) and of a higher molecular weight dye (Texas Red; 70kDa; Invitrogen) was imaged simultaneously at 940nm. Imaging session took place at baseline (before treatment), one day following the first CpG-C/control treatment, and one day following the last treatment. To this end, ten minutes following dye injection, 100μm stacks were taken every ten minutes for a total of ninety minutes. For quantification of dye leakage max-projections of each session were aligned using Fiji software (2.0) plugin *linear stack alignment with SIFT*. Eight vessels (four capillaries 5μm and smaller, and four vessels 20-50μm in diameter) were blindly selected manually and average intensity was measured inside the vessel and adjacent to it (in the parenchyma). The ratio over time between the amount of dye inside and outside the vessels was computed (Fig. 4f). For display purposes only, image contrast was automatically adjusted using the Fiji *autoadjust* display function while measurements were taken directly from pixel values.

### Two-photon laser photodamage

In order to assess microglia reactivity, focal laser-induced thermal damage insults were performed as previously described (44) (Supplementary Fig. 7b). Briefly, CX3CR1^GFP/+^ mice underwent craniotomy and three weeks later microglia were imaged at 890nm. A baseline stack (0-30μm depth) was imaged and a small (~15-20μm) localized injury was achieved by focusing a two-photon laser beam (780 nm; 150mW at the sample; ~1μm size) at 15μm depth for 2s. Stacks were imaged every two minutes for 40min. Using Fiji software (1.0), maximum z-projections were turned into binary images. A 60μm circle was drawn around the ablation area, and, for each time point, number of white pixels were counted inside the small circle (x(t)). For the baseline image, another 120μm circle was drawn, and the number of white pixels in the ring between the two circles were counted (y(0)). Response was calculated as: *x*(*t*) − *x*(0)/*y*(0).

### ImageStream

Preparation of tissue for ImageStream (MK II; Amnis) FACS analysis – mice were perfused with PBS supplemented with 30U heparin (Sigma-Aldrich), and brains removed. Brains were mechanically minced, suspended in a solution containing collagenase (0.1%w/v; Worthington) and dispase (0.2%w/v; Roche) for 20 minutes and then in DNAse (Sigma-Aldrich) for 20 minutes, and suspensions were passed through a 70μm filter. Fatty tissue was removed using Percoll (Sigma-Aldrich), and cells were re-suspended in PBS supplemented with 1% EDTA (Sigma-Aldrich), 0.01% NaN_3_ (Sigma-Aldrich), and 1% FBS (Biological Industries). In each experiment 1×10^4^ events were collected and analyzed using Amnis IDEAS software (Version 6.2). Analysis gates were manually corrected based on images of the events. Internalization was quantified automatically using the software’s internalization wizard (Fig. 6f-h, 7b; Supplementary Fig. 8).

For quantification of CpG-C infiltration into the brain and its internalization by endothelial cells, astrocytes, and microglia (Fig. 3), mice were injected with FITC-labeled CpG-C 24 hours before sacrifice, and single cells suspensions were stained using anti-CD31 (PCAM-1) PE-Cy7 (eBioscience), Anti-GLAST (ACSA-1)-PE (MACS), and anti-CD11b APC (BioGems). To avoid GLAST staining of Bermann glia, cerebellums were removed before preparation of the samples in this experiment. In each population, we quantified the percent of cells with internalized FITC.

**Figure 3:**
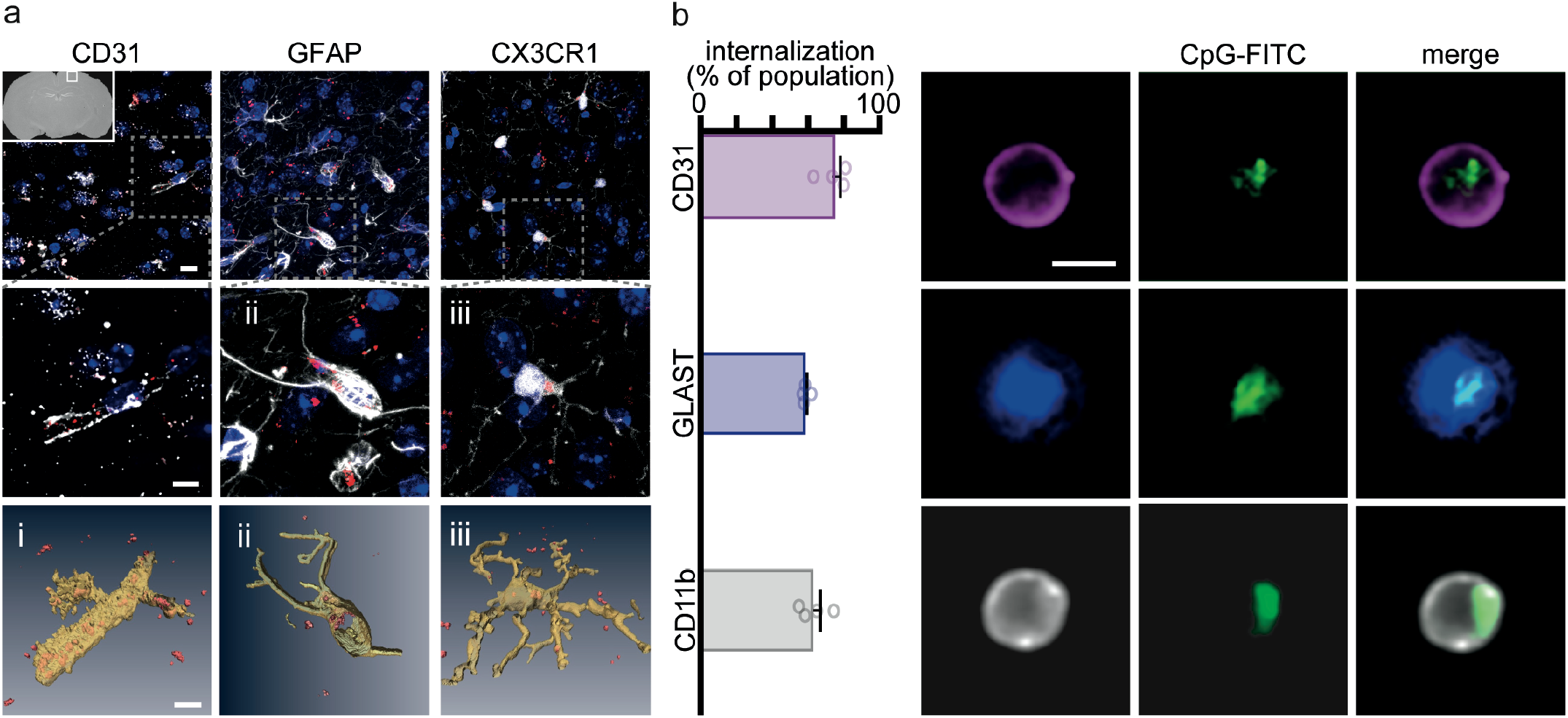
CpG-C infiltrates the brain and is internalized by endothelial cells, astrocytes, and microglia. (**a-b**) TAMRA-labeled CpG-C was injected intraperitoneally, 24 hours later brains were perfused, and CpG-C internalization in endothelial cells (CD31), astrocytes (GFAP and GLAST), and microglia (CX3CR1 and CD11b) was visualized in histological sections using confocal microscopy (**a**; top panels are 15μm z-max projections, and lower panels are partial reconstruction); and quantified using ImageStream FACS analysis (**b**). The majority of each of the three cell populations internalized CpG-C, indicating that CpG-C crosses the BBB into the parenchyma (n=4). Scale bar is 5μm. Data presented as mean (±SEM).

### Quantitative polymerase chain reaction

In two independent experiments (Fig. 8), male and female mature (4-6 months) CX3CR1^GFP/+^ mice were treated with CpG-C or non-CpG ODN/PBS. Twenty-four hours later mice were perfused, brains were harvested and processed into a single-cell suspension as described above. GFP-positive cells (microglia) were sorted (FACSAria IIU, BD Biosciences), and RNA was extracted with TRIzol^®^ (Invitrogen). cDNA was prepared and used for quantitative PCR and the results were normalized to *Gapdh*. All primers and probes were purchased from Applied Biosystems, *Cd36* (Mm00432403_m1), *Cd47* (Mm00495006_m1), *Cd68* (Mm03047343_m1), *Fasl* (Mm00438864_m1), *Gapdh* (Mm99999915_g1), *Inf-γ* (Mm01168134_m1), *Il1-β* (Mm00434228_m1), *Il-6* (*Mm00446190_m1*), *Marco* (Mm00440250_m1), *Nos2* (*Mm00440502_m1*), *Tmem119* (Mm00525305_m1), *Tnf* (Mm00443258_m1), *Tnfsf10* (Mm01283606_m1), *Trem2* (Mm04209424_g1). One control sample was removed as an outlier from statistical analysis of *Tnf* and *Inf-γ* (25 SEMs and 50 SEMs, respectively).

### Volumetric image display

Three-dimensional volumetric reconstruction of single cells (Fig. 3a, Supplementary Fig. 5a) or fields of view (Fig. 6c) where performed in a semi-automatic way using Amira software (ThermoFisher Scientific). Auto-thresholding mode was initially used to detect the brightest object, which depending on the experiment and the spectral channel under analysis, represented either cell soma or aggregates of labeled CpG-C. Cell morphology was partially reconstructed by manual labeling after thresholding.

### Statistical analysis

Prism (version 7.0c) and Python (version 3.6.3) were used for statistical analysis. Where appropriate, the Kolmogorov–Smirnov normality test was used to determine normal distribution of the data, and the F-test or Brown-Forsythe tests for determining homogeneity of variance. For normally distributed data with equal variance, we used one-way ANOVA (Figs. 5a-g, 7b, Supplementary Figs. 6a,b), two-way ANOVA (Figs. 1c.i,e.i, 3b, 6e, Supplementary Figs. 2e, 4, 7), two-tailed unpaired Student’s t-test (Figs. 1d,e.ii,g, 4a-c, 5h,i, 6h, Supplementary Fig. 1d), or one-tailed unpaired Student’s t-test (Fig. 8) to compare experimental groups. For normally distributed data with unequal variance we used Mann–Whitney U-test (Figs. 1c.ii,f, 6f,g Supplementary Fig. 1c, 2a,b) or Kruskal-Wallis (Supplementary Fig. 2c,d) to compare experimental groups. For non-normally distributed data we used two-way permutations (Figs. 2, 7a,c) to compare experimental groups. For post-hoc analysis, multiple comparisons were corrected using Dunn’s test, Tukey’s, or Bonferroni’s, according to the primary analysis and the software’s recommendation. For quantification of primary tumor growth dynamics (Supplementary Fig. 1b) and for longitudinal BBB leakage (Fig. 4e,f), we applied a least squares fit model of an exponential growth curve or one-phase exponential decay curve, respectively, and compared fits of the treated and control groups. p-values smaller than 5% were considered significant. In all experiments, measurements were taken from distinct samples (different animals for *in vivo* experiments and different wells for *in vitro* experiments).

**Figure 4:**
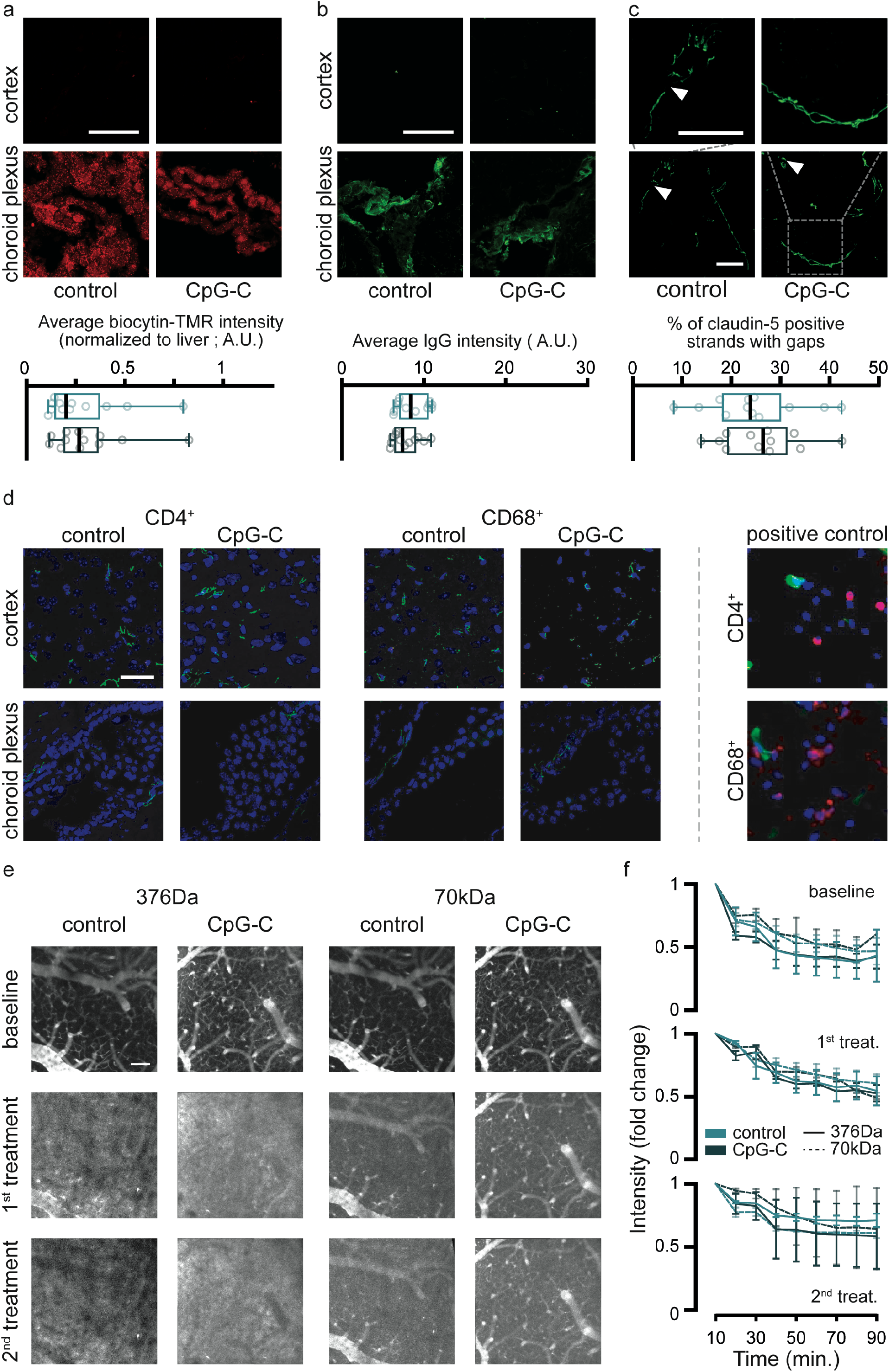
CpG-C does not affect BBB leakage or cellular permeability. (**a-c**) Biocytin-TMR was intravenously injected to animals expressing GFP under the Claudin-5 promotor 24 hours following CpG-C or control treatment, and ninety minutes later brains were perfused and removed. Biocytin-TMR intensity (normalized to intensity levels in the liver, not shown) and IgG staining intensity were similar in CpG-C-treated and control animals (two-tailed unpaired student t-test, t(22)=0.3758, p=0.7106; **a-b**). Moreover, no difference in number of gaps in claudin-5 strands was found between control and CpG-C-treated animals (two-tailed unpaired student t-test, t(22)=0.4283, p=0.6726; **c**). (**d**) Brain sections of WT animals treated with CpG-C or PBS were stained for CD4 or CD68 twenty-four hours following treatment. No infiltration of immune cells was detected (spinal cords of EAE mice served as positive controls; right panels). (**e-f**) Using two-photon imaging we followed leakage of a low (NaF; 376Da) and a high (Texas-Red; 70kDa) molecular weight dextrans. Intensities of representative images (**e**) were auto-adjusted in Fiji for display purposes only. No differences were found between control and CpG-C treated animals at baseline (p=0.8567 and p=0.8421 for NaF and Texas-Red respectively), following a single CpG-C treatment (p=0.9243 and p=0.2419 for NaF and Texas-Red respectively), and following two CpG-C treatments (p=0.4656 and p=0.3918 for NaF and Texas-Red respectively. See methods for an explanation of the quantification; **f**). For (**a-c**), each sample consisted of an average of at least five images that were analyzed. Samples were taken from four different anatomical brain regions (cortex, midbrain, cerebellum, and hippocampus) in three mice/group (See Supplementary Fig. 4 for regional presentations). Scale bar is 50μm. Boxplot whiskers represent min-max range (**a-c**) and data in (**f**) is presented as mean (±SEM).

**Figure 5:**
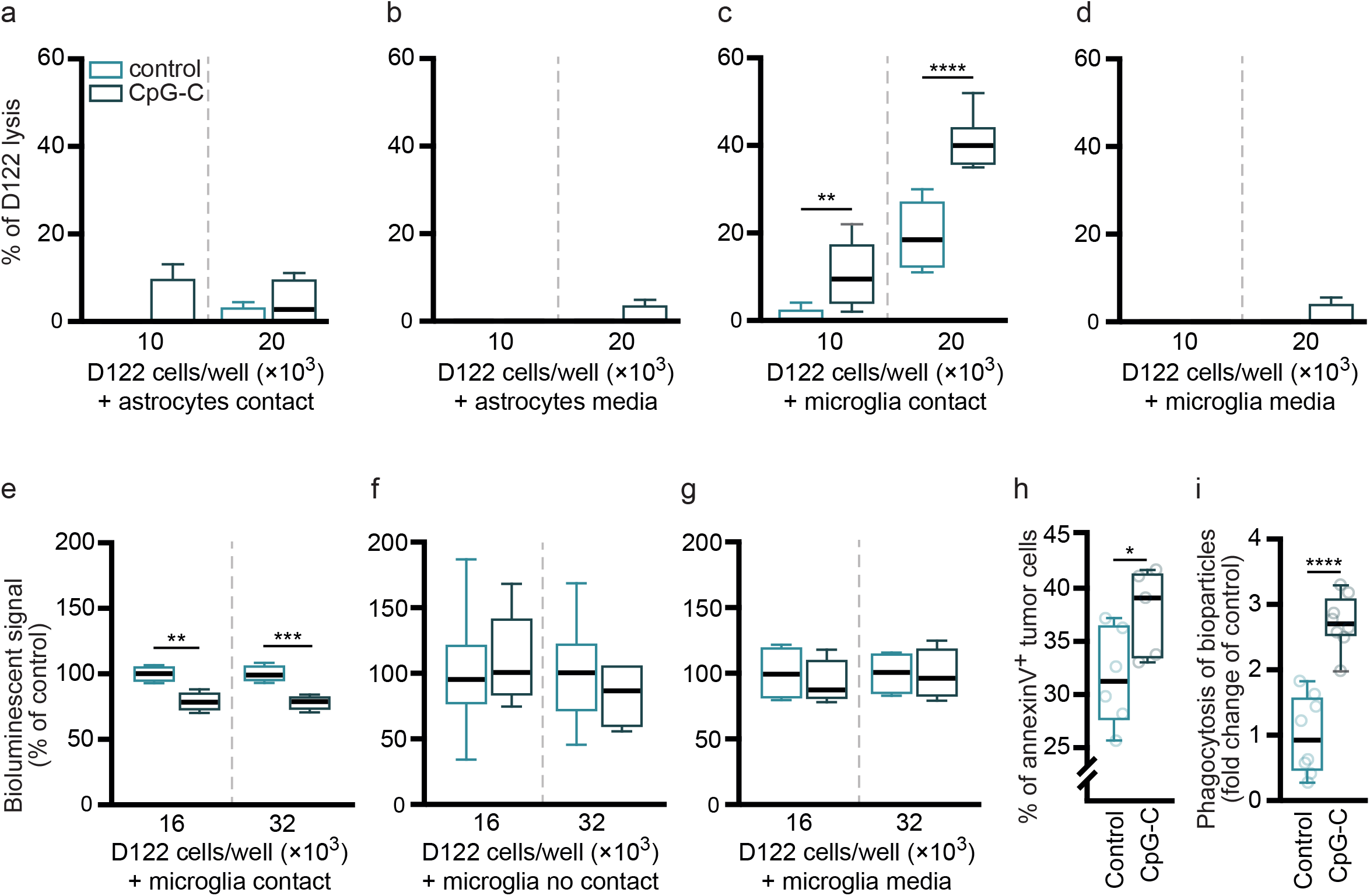
Microglia, but not astrocytes, mediate the effects of CpG-C *in vitro*. (**a-d**) Primary cultures of microglia and of astrocytes were subjected to 100nM/L CpG-C or non-CpG ODN (control) for 24 hours. ^125^IUDR-labeled D122 cells were plated with the treated primary cultures, or subjected to their conditioned-media alone, and cytotoxicity (percent of D122 lysis) was assessed by measuring radioactivity in the media 8 hours later. Primary cultured astrocytes, subjected to CpG-C or non-CpG ODN, did not cause tumor cell lysis when in contact (F_(3,12)_=0.7755, p=0.5298; **a**), nor did their conditioned-media (F_(3,9)_=0.6923, p=0.5794; **b**). In contrast, primary cultured microglia cells induced lysis in tumor cells, and treatment with CpG-C significantly increased it when in contact with tumor cells (F_(3,28)_=64.1, p<0.0001; **c**), while their conditioned-media had no effect (F_(3,9)_=0.6923, p=0.5794; **d**). (**e-i**) The microglia cell line, N9, was subjected to CpG-C (see above). Luc2-mCherry-labeled D122 cells were plated with the N9 cultures with contact (**e**) or without contact (co-culture; **f**), or with their conditioned-media alone (**g**), and bioluminescent signal was measured, indicating viability of tumor cells. There was a reduced signal in tumor cells cultures that were in direct contact with N9 cells (F_(3,12)_=14.6, p=0.0003; **e**), while no difference was evident in co-cultures (no contact; F_(3,18)_=0.3535, p=0.7872; **f**), or in cultures subjected to conditioned-media (F_(3,12)_=0.1425, p=0.9325; **g**). Two-tailed one-way ANOVA with Bonferroni’s multiple comparison correction was used for **a-g**. (**h**) Annexin V binding (a marker for early stage apoptosis) was quantified in D122 cells cultured with pretreated N9 cells using FACS. Tumor cells cultured with N9 cells pretreated with CpG-C exhibited increased annexin V staining (compared to scrambled CpG-C; two-tailed student’s t-test for unpaired samples, t(9)=2.306, p=0.0465). (**i**) N9 cultures treated with CpG-C or non-CpG ODN (control) for 24 hours were washed and plated with pH-sensitive bio-particles to assess phagocytosis capacity. CpG-C treated N9 cells exhibited a 3-fold increased phagocytic capacity (two-tailed student’s t-test for unpaired samples, t(14)=6.696, p<0.0001). Boxplot whiskers represent min-max range. Refer to supplementary figure 6b for comparison between PBS, non-CpG ODN, and CpG-C.

## RESULTS

### Prophylactic CpG-C treatment reduces brain metastases in experimental and spontaneous metastases models

To study the prophylactic efficacy of the TLR9 agonist CpG-C in reducing brain metastasis, we first employed two models of non-small-cell lung carcinoma, given the clinical prevalence of brain metastases in this type of cancer (15,45). To this end, we used the highly metastatic D122 variant of the syngeneic Lewis lung carcinoma (LLC) in C57BL/6 mice (32), and the human xenograft PC14-PE6 cells in athymic nude mice (46). For exclusive injection of tumor cells to the cerebral circulation, we used a novel approach that we have recently developed and validated – the assisted external carotid artery inoculation (aECAi; Fig. 1a,b) (32) – which results in improved targeting of tumor cells to the brain, and avoids cerebral blood flow perturbations. A single prophylactic systemic injection of CpG-C was given 24 hours before tumor cell injection. Brain tumor growth was monitored thereafter using *in vivo* bioluminescence imaging. Animals pretreated with CpG-C displayed reduced cerebral tumor growth in both the syngeneic (p=0.0011; Fig. 1c) and the xenograft (p<0.0001; Fig. 1e) models, exhibiting a statistically significant difference starting on days 14 and 4, respectively. At end point, signal intensity, which is indicative to the tumor burden, was 77-fold lower in the CpG-C treated mice in the syngeneic D122 model (n=6; p<0.0001), and 82-fold lower in the xenograft model (n=7; p<0.0001), compared to their matching control groups (n=7 in both models). To assure that the differences between groups originated from tumor growth within the brains, rather than from extra-cranial growth (32), we harvested the brains and measured the tumor signal in both models. In the syngeneic model, brains from CpG-C treated animals had 48-fold lower bioluminescent signal compared to control animals (p=0.0040; Fig. 1d). Similarly, in the xenograft model the mean area of fluorescence signal, indicative of brain tumor burden, was significantly smaller in CpG-C treated animals (p=0.0373; Fig. 1f).

To test the efficacy of CpG-C treatment in a context that better resembles the clinical setting, we used a murine model of spontaneous brain metastasis that we have recently established (33). In this model, mCherry-expressing Ret melanoma cells are injected orthotopically, resulting in growth of a primary tumor in the flank. During the perioperative period – three days before and after primary tumor excision – animals were treated with CpG-C (n=15) or vehicle (n=11), with no measurable impact on primary tumor growth (p=0.6066 for tumor growth dynamics and p=0.9260 for tumor size at time of excision; Supplementary Fig. 1a-c). Approximately nine weeks after excision of the primary tumor, brain and lung metastatic burden (i.e. mCherry expression) were quantified. CpG-C treatment significantly reduced the overall metastatic burden in the brain (n=9 and 12 for control and CpG-C, respectively; p=0.0345; Fig. 1g). Notably, in the lungs (as in the primary tumor) CpG-C treatment had no effect (p=0.7858; Supplementary Fig. 1d), suggesting that the beneficial effects of CpG-C in the brain were not secondary to generic or peripheral effects on tumor burden. These results provide direct evidence that systemic prophylactic CpG-C treatment during the perioperative period can reduce metastatic growth in the brain.

### Prophylactic CpG-C is effective in reducing tumor seeding in the brain in a variety of treatment settings

In the subsequent experiments, we aimed to pinpoint mechanisms underlying the beneficial effects of CpG-C. We focused on the first 24 hours of tumor colonization in the brain, for the following reasons: (i) a single administration of CpG-C, that we herein found effective, is known to exert immune activation within hours and for up to 72 hours (11); (ii) bioluminescence imaging indicated a non-significant trend for beneficial effects a day following tumor inoculation (data not shown); and (iii) tumor cells successfully proliferate to macrometastases only if they extravasate into the brain parenchyma within the first three days (46). To maximize our ability to focus on the first days following CpG-C administration, we administered syngeneic D122 tumor cells in C57BL/6J mice, employing the aECAi approach (32), known for its high temporal inoculation efficiency. We assessed brain tumor seeding, measuring radioactive signals of isotope-labeled tumor cells within an entire excised organ – an approach that allows maximal signal-to-noise sensitivity.

A single prophylactic injection of CpG-C resulted in reduced brain tumor retention, similarly in males and females (Supplementary Fig. 2a), and young, juvenile, and old mice (Supplementary Fig. 2b). While CpG-C was effective in reducing brain tumor retention already at a dose of 1.2mg/kg (p=0.0455), its efficacy increased at 4mg/kg (p=0.0003; Supplementary Fig. 2c) – a dose we previously showed as beneficial in reducing peripheral metastases (39). In the clinical setting, a prophylactic treatment should rely on a chronic schedule, and therefore, we tested whether a regime of five injections of CpG-C given every other day has similar effects as a single injection, and does not result in tolerance to the effects of the agent. Indeed, CpG-C treatment resulted in reduced tumor retention (p=0.0001; Supplementary Fig. 2d), following both the acute (n=6; p=0.0298) and the chronic (n=6; p=0.0013) treatments, compared to control animals (n=6). Notably, single and multiple CpG-C injections were well tolerated, as indicated by a lack of weight loss compared to control animals (n=6; p=0.2593; Supplementary Fig. 2e), in line with previous reports (11). These data suggest that CpG-C is efficient both as an acute and as a chronic prophylactic treatment for brain metastasis, in both sexes and across ages, affecting early stages of tumor cell seeding.

### NK cells and macrophages are not involved in the metastatic process in the brain, nor mediate the beneficial effects of CpG-C

It has previously been shown that CpG-ODNs have beneficial effects in the periphery, reducing seeding of tumor cells, and their subsequent growth. These anti-tumor effects were found to be mediated by NK cells (12,47) and macrophages (10). To study *in vivo* whether these leukocytes also take part in the metastatic process in the brain and mediate the effects of CpG-C, we depleted NK cells and monocytes/macrophages using anti-NK1.1 and clodronate liposomes, respectively (Fig. 2). In the lungs, NK depletion resulted in a 5-fold increase in tumor retention (p=0.0001), and partially blocked the beneficial effects of CpG-C (p=0.0038; Fig. 2a) evident in naïve animals (p=0.0019) (in line with previous results (48)). In contrast, in the brains of the same animals NK depletion did not affect tumor retention (p=0.3935), nor did it mediate the beneficial effects of CpG-C (p=0.0811; Fig. 2b), evident in both naïve (p<0.0001) and NK-depleted animals (p=0.0056). Similarly, depletion of monocytes increased lung (p=0.0401; Fig. 2c), but not brain tumor retention (p=0.3081; Fig. 2d), while the effects of CpG-C were not mediated by monocytes in the lungs (p=0.0003) or in the brain (p=0.0001). These findings demonstrate that NK cells and monocytes/macrophages play a key role in the metastatic process in the lungs, but not in the brain, nor do they mediate the beneficial effects of CpG-C in the brain.

### CpG-C is taken up by cerebral cells without disrupting blood-brain barrier integrity

As peripheral innate immune cells do not seem to mediate the effects of CpG-C, we turned to evaluate the role of CNS cells that express TLR9 (23,49). We focused on cells that are known to play key roles in the metastatic process, including endothelial cells, astrocytes, and microglia (50). First, to evaluate whether CpG-C can cross the BBB and affect cerebral components, we systemically administered mice with TAMRA- or FITC-conjugated CpG-C. Twenty-four hours later we analyzed CpG-C uptake by brain endothelia, astrocytes, and microglia, in histological sections (Fig. 3a) and using ImageStream FACS analysis (see methods; Fig. 3b). Approximately 74% of endothelial cells, 58% of astrocytes, and 62% of microglia cells internalized CpG-C (n=4 mice; Fig. 3b). This internalization is expected, as TLR9 ligands are internalized into the cell to bind with the endosomal receptors (51). Indeed, lysosomal staining of microglia extracted from CpG-C-TAMRA treated animals indicated CpG-C is internalized into the lysosomes (Supplementary Fig. 5b).

For malignant cells to infiltrate into the brain parenchyma, they must cross the BBB. Endothelial cells connected by tight junction act as the first physical barrier, preventing uncontrolled infiltration of blood-borne cells. As endothelial cells uptake CpG-C (Fig. 3a,b), we sought to test whether it had an effect on BBB permeability and integrity. To this end, we measured Biocytin-TMR and IgG infiltration and continuity of Claudin-5 (tight junctions) in animals expressing GFP under the Claudin-5 promotor. CpG-C did not affect Biocytin-TMR of IgG infiltration (≥5 images were averaged in 4 anatomical regions in 3 mice – n=12; Fig. 4a,b Supplementary Fig. 4a-c), nor continuity of Claudin-5 (Fig. 4c, Supplementary Fig. 4d). Furthermore, no infiltration of immune cells (i.e. CD4^+^ or CD68^+^) was evident following CpG-C treatment (Fig. 4d). Thus, these results strongly suggest that the effects of CpG-C on tumor seeding in the brain are not mediated by perturbations to the BBB or the choroid plexus.

### Microglia, but not astrocytes, mediate anti-tumor beneficial effects of CpG-C

Astrocytes (52) and microglia (53) have key roles in innate and adaptive immunity, and combined with their significant uptake of CpG-C (Fig. 3ab and Supplementary Fig. 5a), they were our primary candidates for mediating the effects of this agent. Therefore, we investigated their *in vitro* capacity to induce tumor cell lysis and the impact of pre-stimulation with CpG-C. Primary astrocytic cultures were treated with CpG-C or non-CpG ODN, and tested for their ability to induce tumor cell lysis by contact or by secretion of apoptosis-inducing factors. The cultured astrocytes did not induce tumor cell death, with or without CpG-C treatment, in both contact and secretion conditions (Fig. 5a,b). In contrast, primary microglial cells induced cytotoxicity in D122 tumor cells, and CpG-C treatment markedly increased this lysis when tumor cells were in contact (Fig. 5c), while their conditioned-media alone had no effect (Fig. 5d). We further extended this testing in the N9 immortalized microglia cell line. Similar to the effects observed in the primary microglia culture, N9 cells reduced tumor cell viability when in contact (Fig. 5e), but failed to do so in a paracrine setting (Fig. 5f,g). To study whether non-CpG ODN impacted the tumoricidal activity of N9 cells, we repeated the contact co-culture experiment with an additional group of PBS-treated N9 cultures (Supplementary Fig. 6b). We found PBS and non-CpG ODN treatments to have a similar affect (p=0.7745 and p=0.1420 for 16▯10^3^ and 32▯10^3^ D122 cells/well), while CpG-C significantly reduced tumor cells viability (for 16▯10^3^: p=0.0017 and p=0.0062 compared to PBS and non-CpG ODN, respectively, and for 32▯10^3^: p=0.0003 and p=0.0477 compared to PBS and non-CpG ODN, respectively). Next, we studied the mechanisms by which microglia cells eradicate D122 tumor cells. We found that N9 cells treated with CpG-C induced apoptosis in tumor cells, as indicated by increased annexin V staining (Fig. 5h). Additionally, CpG-C treatment resulted in a 3-fold elevation in phagocytosis capacity (Fig. 5i), in line with previous reports (27). Notably, it appears that the effect of CpG-C on microglia activity is not a general activation, as we found no effects of the agent in a scratch migration assay (54) (n=9; p=0.6732 for wound confluency, and p=0.6039 for wound width; and also *in vivo* as described below, Supplementary Fig. 7a). Taken together, these findings indicate that contact between microglia and tumor cells is essential for the effects induced by CpG-C. A combination of elevated microglial cytotoxicity and enhanced phagocytic capacity underline these effects.

### Microglia mediate the beneficial effects of CpG-C *in vivo*

We found that CpG-C affects brain tumor retention as early as 24 hours post tumor cell inoculation. Interactions between microglia and tumor cells at early stages of tumor cell extravasation have been reported elsewhere (55). However, the significance of these interactions with respect to microglial tumoricidal characteristics at this time point, is yet unknown. To this end, we first established that microglia indeed phagocytize tumor cells at this early time point. Longitudinal intravital imaging revealed that microglia cells interact with tumor cells, and initiate phagocytic processes, as early as a few hours after tumor cell inoculation (Fig. 6a-c). To assess the effects of CpG-C on this phagocytic capacity, CX3CR1^GFP/+^ mice were injected with CpG-C or CpG non-ODN, and 24 hours later injected with either tdTomato-labeled or mCherry-labeled D122 tumor cells for two-photon or ImageStream FACS analysis, respectively. The number of microglia-tumor cells contacts and microglia internalization of mCherry particles (originated from tumor cells) were quantified four hours following tumor cells inoculation and at days one, four, and seven thereafter (Fig. 6d,e). As early as four hours following tumor cells inoculation there were more contacts between microglia and tumor cells in CpG-C treated animals (p=0.0128), with no differences at later times. Moreover, the number of internalization events CpG-C treated animals were higher four hours (p=0.0372) and one day (p=0.0041) following tumor cell inoculation. No differences were evident at days four and seven, probably due to the dismantling process of the tumor cells evident as early as two days following tumor cell inoculation (Fig. 6a). Using ImageStream FACS analysis 24 hours after tumor inoculation, we found first that CpG-C did not affect the total number of microglia in the brain (n=5; p=0.4201; Fig. 6f), nor the total number of infiltrating tumor cells (p=0.3455; Fig. 6g), in accordance with our above findings regarding the lack of CpG-C impact on BBB permeability. However, CpG-C increased phagocytosis of tumor cells by microglia (p=0.0055; Fig. 6h). These results alone do not specify whether CpG-C increases the killing of tumor cells by microglia, or whether it merely increases endocytosis of tumor debris by microglia. To distinguish between these alternatives, we turned to a set of experiments where microglia activation was impaired or where microglia were depleted from the brain and quantified the ability of CpG-C to reduce the total amount of live tumor cells, by assessing radioactive signaling that originated from radio-labeled tumor cells. Employing this approach, animals were treated with minocycline, an inhibitor of microglial activation (40,56) (Fig. 7a), which resulted in a significantly increased brain tumor retention (p=0.0118), without affecting the total number of infiltrating tumor cells (see below). Importantly, CpG-C treatment reduced tumor retention in naïve mice (p<0.0001), but not in minocycline-treated animals (p=0.1863). Moreover, the effects of CpG-C were completely blocked by minocycline treatment (p<0.0001), indicating the mediating role of microglia in the beneficial effects of CpG-C. To further validate these significant results, animals were treated with CpG-C, or with minocycline and CpG-C, and mCherry (tumor cells) uptake by microglia was quantified using ImageStream FACS analysis, and compared to saline-treated animals (Fig. 7b). In line with the radioactive-based quantification, CpG-C increased tumor cell phagocytosis (i.e. events where the mCherry signal could be identified inside GFP-positive segmented objects; p=0.0100), and this effect was blocked by minocycline (p=0.0493). Notably, infiltration capacity of tumor cells was not affected by minocycline treatment, as indicated by total area of mCherry (i.e. all detection events combined) in the brain (p=0.8994). Depletion of all microglia (activated and non-activated) with PLX5622 (41), a colony-stimulating factor 1 receptor (CSF1R) inhibitor, blocked the beneficial effects of CpG-C (p=0.0068), again indicating the mediating role of microglia. Microglia depletion alone did not affect tumor retention in brains of naïve animals (p=0.7490; Fig. 7c).

**Figure 6:**
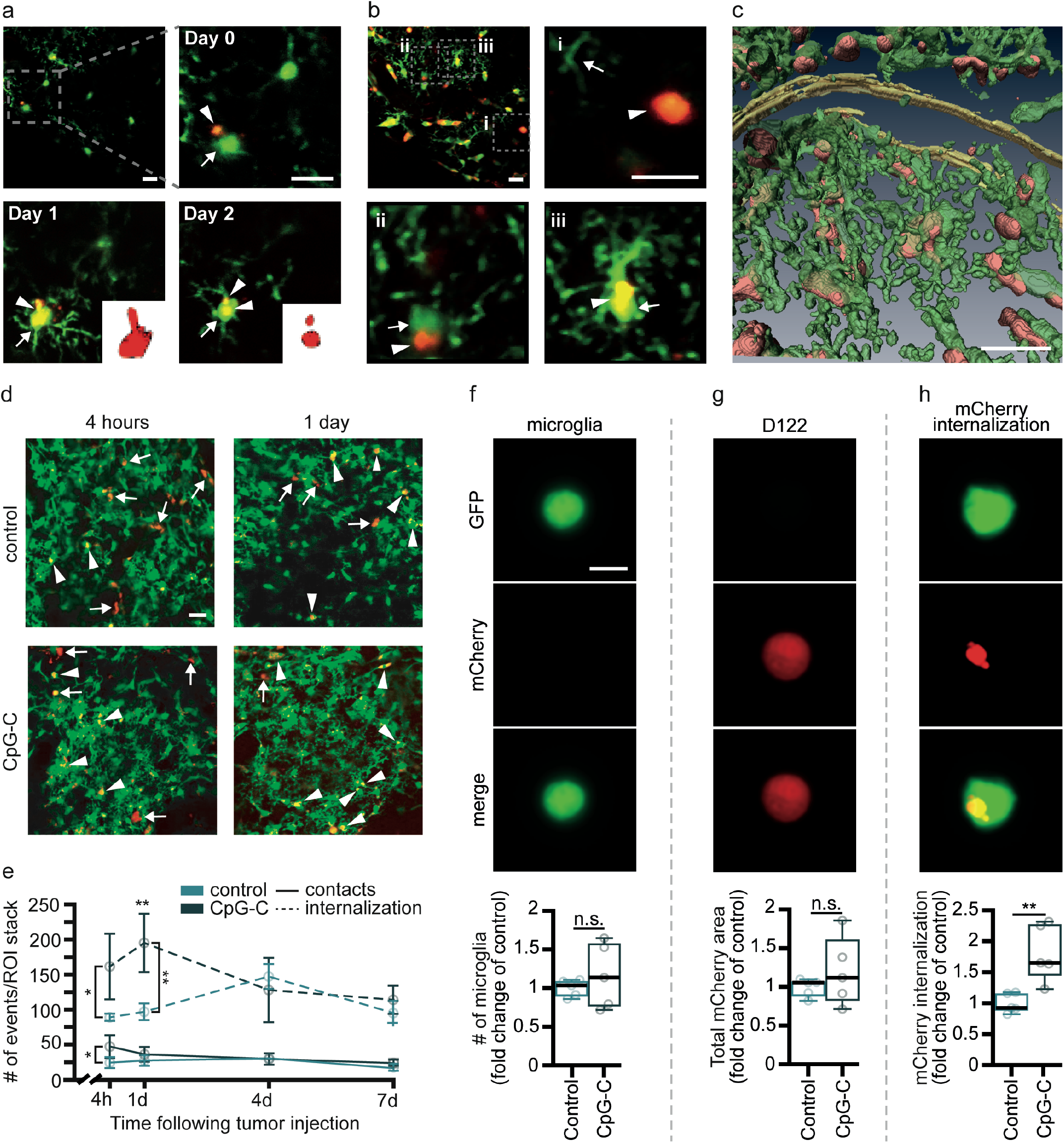
Microglia mediate the *in vivo* effects of CpG-C. (**a-e**) Chronic *in vivo* two-photon imaging in CX3CR1^GFP/+^ mice indicated microglia cells (green) have dynamic relations with tdTomato-labeled D122 tumor cells (red; 15μm stacks, with 1μm z-steps) and that CpG-C treatment increases tumor internalization by microglia. (**a**) A microglia cell (arrow) interacting with a tumor cell (arrow head; two hours post tumor cells inoculation), phagocytizing it (one day later), and dismantling it (day 2 post tumor cell injection, inset shows an engulfed tdTomato-positive cell or part of it). (**b**) Different levels of interaction between microglia and tumor cells - (i) no interaction; (ii) contact; and (iii) microglia phagocytized a tumor cell. (**c**) Partial reconstitution of a 15 μm stack with 1μm z-steps demonstrating the microglia-tumor cells’ “battle field” four hours after tumor cell injection. (**d**) Representative images and quantification (**e**) of microglia-tumor interactions in control and CpG-C treated mice four hours and one day following tumor cell inoculation (arrows for contact and arrow heads for internalization). CpG-C treatment resulted in increased contacts four hours following tumor inoculation (n=3; F_(1,4)_=2.875, p=0.0218) and in microglia internalization of tumor cells/debris four (p=0.0372) and twenty-four hours (n=3; F_(1,4)_=3.400, p=0.0041) following tumor inoculation. Scale bar for (**a-d**) is 20μm. (**f-h**) CX3CR1^GFP/+^ mice were treated with a single systemic prophylactic CpG-C treatment, injected mCherry labeled D122 tumor cells using the aECAi approach, and brains were analyzed using ImageStream FACS. While CpG-C treatment did not affect the number of microglia cells (n=5; two-tailed Mann-Whitney U=10, p=0.6905; **f**), or capacity of tumor cell infiltration (indicated by total mCherry area in perfused brains; two-tailed Mann-Whitney U=9, p=0.5476; **g**), it resulted in increased phagocytosis of tumor cells by microglia (two-tailed student’s t-test for unpaired samples, t(4)=3.885, p=0.0178; **h**). Scale bar for (**e-g**) is 5μm. Data in (**e**) is presented as mean (±SEM) and boxplot whiskers represent min-max range (**f-h**).

**Figure 7:**
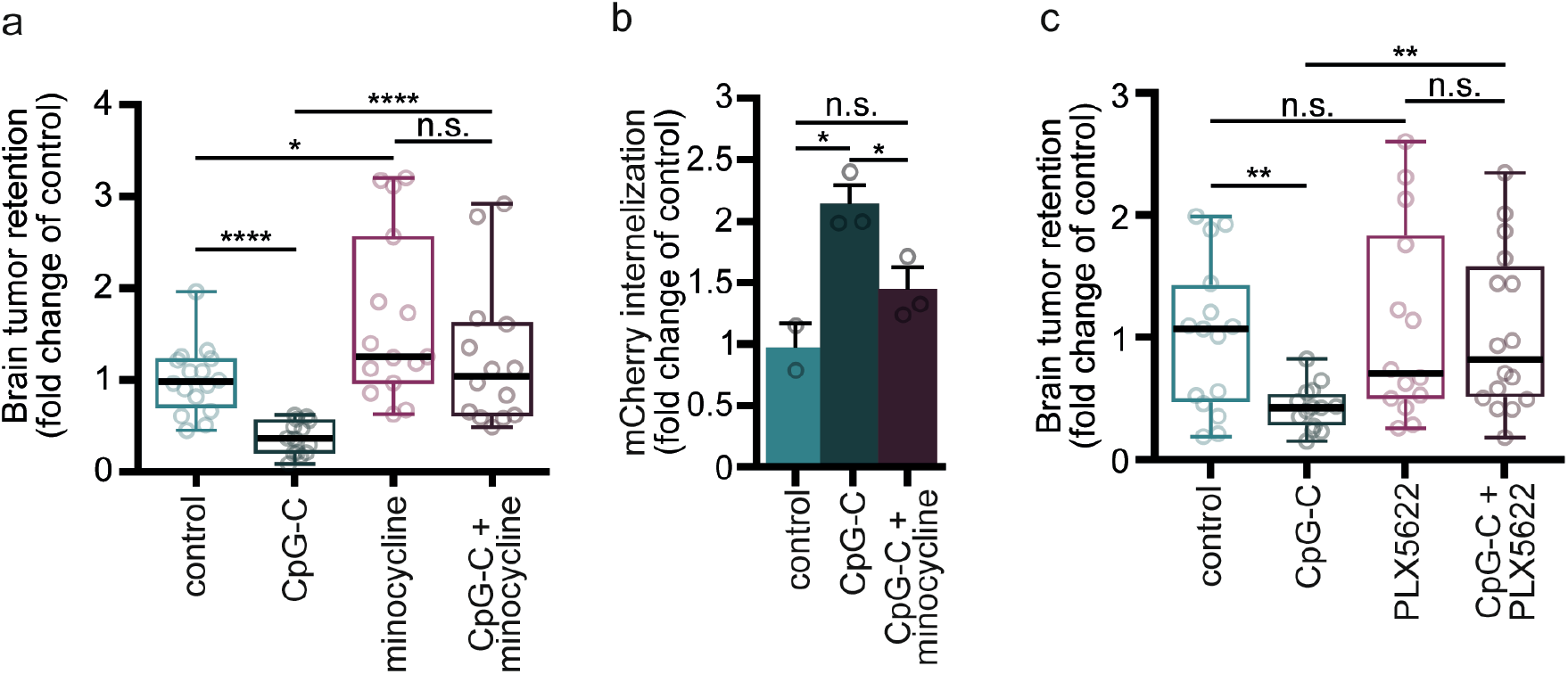
Blocking microglia activation or complete depletion hinders the effects of CpG-C on brain metastasis. (**a-b**) Microglia activation was blocked *in vivo* using systemic treatment with minocycline. (**a**) Minocycline treatment resulted in increased brain tumor retention of D122 cells (n=15-16, t(27)=66.3229, p=0.0118), while CpG-C treatment reduced tumor retention in naïve mice (n=16 for control and n=14 for minocycline treated mice; t(28)=63.0149, p<0.0001), but not in minocycline-treated animals (n=14-15; t(27)=42.7850, p=0.1863). The effects of CpG-C were completely blocked by minocycline treatment (n=14; t(26)=86.5528, p<0.0001), indicating that microglia activation mediates the beneficial effects of CpG-C. (**b**) ImageStream FACS analysis indicated that minocycline blocked (p=0.0493) the beneficial effects of prophylactic CpG-C treatment (p=0.01) on the ability of microglia to phagocytize tumor cells (n=2 for control and n=3 for CpG-C and CpG-C+minocycline animals; two-tailed one-way ANOVA with Tukey’s multiple comparisons test; F_(2,5)_=12.85, p=0.0107). (**c**) Without stimulation with CpG-C, microglia cells do not affect brain tumor seeding, as indicated by depletion of microglia cells using the colony-stimulating factor 1 receptor inhibitor, PLX5622. Microglia depletion did not affect D122 tumor retention in the brain (n=14-15; t(27)=0.0851, p=0.7490), while it blocked the beneficial effects of CpG-C (n=14 for depleted animals and n=16 for depleted animals treated with CpG-C; t(28)=0.0460, p=0.8637), evident in naïve animals (n=14-15; t(27)=0.0460, p=0.0087). Accordingly, microglia-depleted animals treated with CpG-C had increased brain tumor retention compared to naïve animals treated with CpG-C (t(28)=0.5417, p=0.0068). Two-way permutations were used for analyses of (**a**) and (**c**). Boxplot whiskers represent min-max range (**a,c**) and data in (**b**) is presented as mean (±SEM).

Given our *in vitro* and *in vivo* results, we predicted that CpG-C administration would result in elevated expression of apoptosis- and phagocytosis-related factors by microglia cells. We therefore preformed transcriptional analysis of microglia cells isolated from CpG-C treated, or control, animals (Fig 8a). We revealed a robust impact of the agent on the induction of mRNA encoding of apoptosis-inducing, phagocytosis related, and inflammatory factors, while not affecting the inflammation-independent microglial marker *Tmem119* (57) (p=0.7258; Fig. 8a). Specifically, mRNA expression of the key apoptosis-inducing ligands, *Tnfsf10* and *Fasl*, increased by 3-4-fold in microglia from CpG-C-treated animals (p=0.0252 and p=0.0324, respectively; Fig. 8b). In addition, CpG-C treatment resulted in increased expression of receptors related to phagocytosis (58), including, CD47 (p=0.0186) and *Trem2* (p=0.0199), while *Cd36* and *Cd68* mRNA expression levels did not change (p=0.7080 and p=0.9874, respectively; Fig. 8c). *Marco* (macrophage receptor with collagenous structure), another important phagocytosis receptor (59), was not detected in microglia of control animals, yet it was highly expressed in CpG-C-treated animals (p=0.0108; Fig. 8c). While mRNA of the inflammatory cytokines *Il-6* and *Il1-β* was not affected by CpG-C treatment (p=0.9690 and p=0.6772, respectively), *Tnf* and *Inf-γ*, which are known to synergistically induce apoptosis in tumor cells (54), were increased by approximately two- and seven-fold, respectively (p=0.0163 and p=0.0374, respectively; Fig. 8d). mRNA of nitric oxide synthase 2 (*Nos2*), an inflammation-associated enzyme with tumoricidal properties at high concentrations (60), was not detected in control animals, while abundantly expressed in CpG-C treated animals (p=0.0203; Fig. 8d). Irrespectively, and in line with our *in vitro* results, CpG-C did not affect microglia reaction to a non-tumor-related stimulus *in vivo* (i.e. laser induced photodamage; p=0.7474; Supplementary Fig. 7b). Overall, these *in vivo* findings strengthen the notion that prophylactic treatment with CpG-C is beneficial in reducing brain metastasis, by triggering non-activated microglia cells to adopt tumoricidal characteristics.

**Figure 8:**
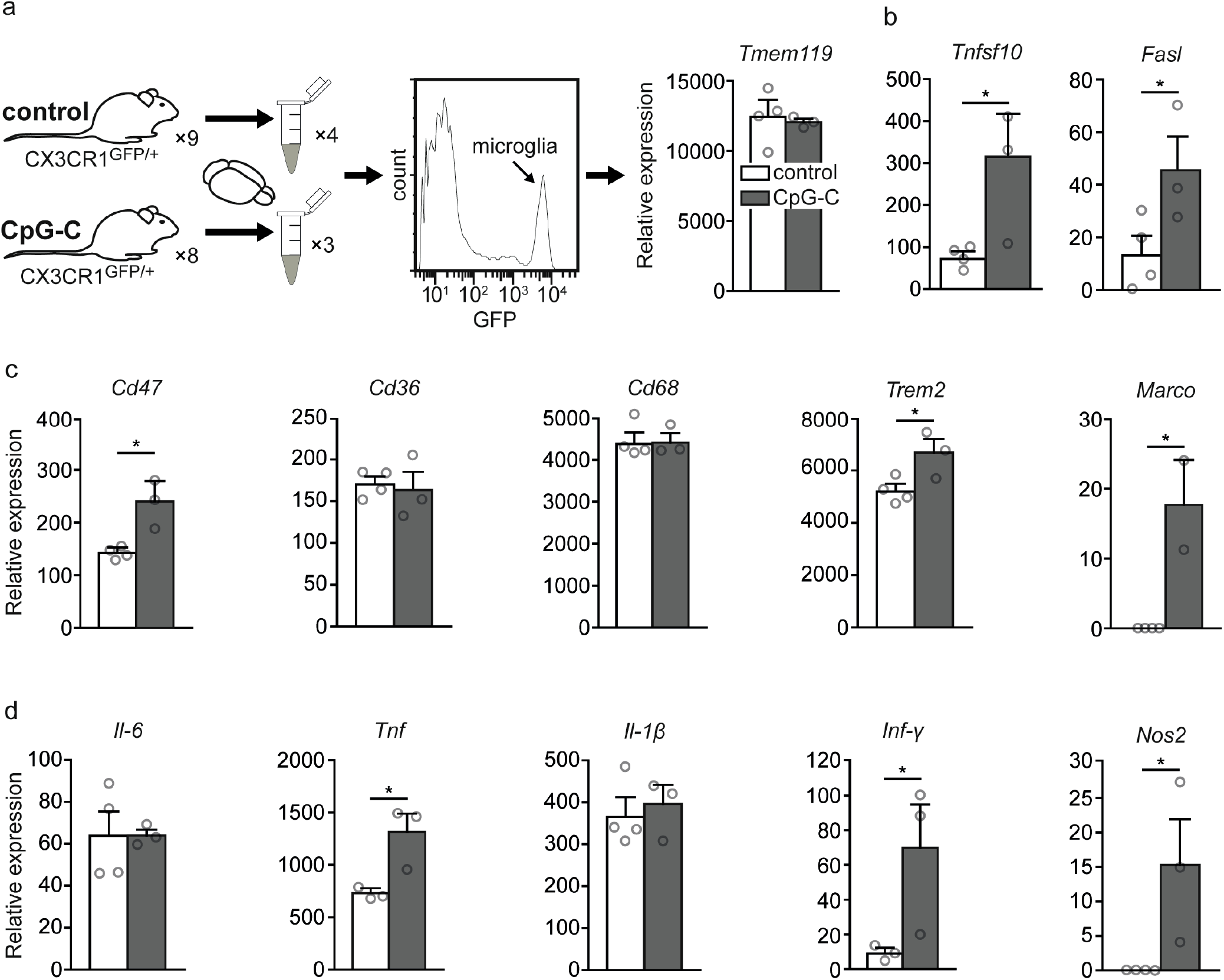
CpG-C treatment results in elevated *in vivo* expression of apoptosis-inducing, phagocytosis-related, and inflammatory factors. (**a**) CX3CR1^GFP/+^ mice were injected with CpG-C or vehicle, and 24 hours later mRNA expression levels in sorted microglia cells were quantified using qRT-PCR. In one experiment six animals of each group were pooled into a single sample, and in the second experiment, two CpG-C treated animals, and three controls, were analyzed separately (n=3-4 from 8-9 animals). As expected, *Tmem119*, a general microglia marker was unaffected by the treatment (t(5)=0.371, p=0.7258). (**b**) The death ligands, *Tnfsf10* and *Fasl*, were elevated by 3-4-fold by a single CpG-C injection (t(5)=2.564, p=0.0437; and t(5)=2.36, p=0.0324, respectively). (**c**) Expression levels of receptors related to phagocytosis were significantly higher in microglia of CpG-C-treated animals. While no change was apparent in *Cd36* (t(5)=0.3966, p=0.7080) and *Cd68* (t(5)=0.01655, p=0.9874), a significant increase was evident in *Cd47* (t(5)=2.819, p=0.0186), *Trem2* (t(5)=2.762, p=0.0199), and *Marco* (which was not detected in control animals) (t(4)=4.499, p=0.0108). (d) While RNA of the inflammatory cytokines *Il-6* and *Il1-β* was not affected by CpG-C treatment (t(5)=0.04089, p=0.9690; t(5)=0.4417, p=0.6772, respectively), *Tnf* (t(4)=3.207), p=0.0163) and *Inf-γ* (t(4)=2.394, p=0.0374), which synergistically induce apoptosis in tumor cells (54), and *Nos2* (t(5)=2.744, p=0.0203), which is tumoricidal at high concentrations (60), were increased following CpG-C injection. Data is presented as mean (±SEM).

**Figure 9:**
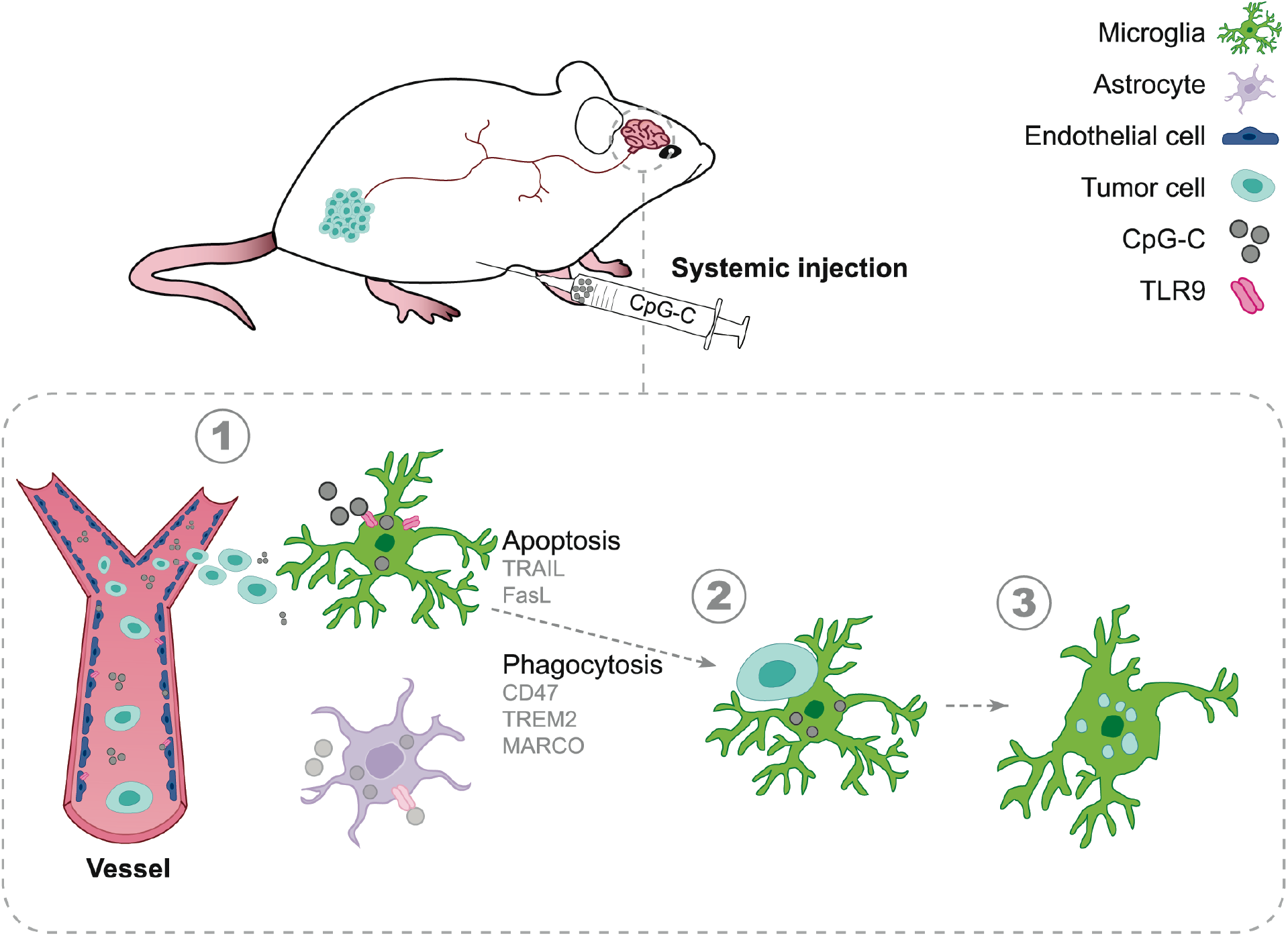
Proposed mechanism. Systemic prophylactic treatment with CpG-C during the perioperative period activates microglia to induce apoptosis in tumor cells and phagocytize them, resulting in reduced brain metastases colonization. A few weeks to months may pass from the time of cancer diagnosis to the time of primary tumor excision (93). During this period, and a few weeks after surgical excision (known as the perioperative period), there is a high risk for developing brain metastasis with terminal consequences (1). CpG-C, a TLR9 agonist, given as a systemic prophylactic treatment during this crucial period, infiltrates the brain and activates microglia (1), increasing their expression of *Tnfsf10* and *Fasl*, resulting in contact-dependent induced-apoptosis of tumor cells (2). Furthermore, *Cd47, Trem2*, and *Marco* expression is increased, triggering enhanced microglial phagocytosis and dismantling of tumor cells (3), thereby reducing brain metastasis colonization.

## Discussion

Brain metastasis is a detrimental manifestation of cancer progression with limited treatments; and a better understanding of this process is expected to improve therapeutic interventions. Here, employing three tumor models, we report that prophylactic systemic treatment with CpG-C, a TLR9 agonist, exerts beneficial effects through reducing tumor cell seeding and growth in the brain. Notably, NK cells and monocytes did not mediate anti-metastatic processes in the brain, nor the beneficial effects of CpG-C, in contrast to their important role in the periphery (shown also here in the lungs). Instead, we identify microglia as key mediators of these beneficial effects in the initial steps of metastatic brain colonization. Moreover, we show that activation of microglia is essential for its anti-metastatic function. Thus, CpG-C stimulates microglia to adopt anti-tumor characteristics, inducing tumor apoptosis and phagocytosis, thereby reducing the formation of brain metastases.

Systemic treatment against brain metastasis has been proposed as a first therapeutic choice (2,4,61), but no effective clinical routine is yet available. A previous study indicated that systemic administration of a CpG-ODN can result in altered cerebral mRNA expression profile (62) suggesting that the agent could have reached this organ. Furthermore, CpG-ODN was shown to stimulate BV2 microglia cells *in vitro* (63), and intracranial injection of CpG-ODN resulted in activation of microglia cells *in vivo* (64). However, there was no direct *in vivo* evidence demonstrating that such an agent could enter the brain parenchyma if administered systemically and elicit a beneficial effect, fundamental requirements for a prophylactic treatment in cancer patients. Notably, direct intracranial injection of tumor cells or CPG-ODN (or any other agent) alter the neuro-immune environment by eliciting an inflammatory response (65), thus, interpreting the role of immune cells in these settings in less straightforward. We overcome these technical hurdles and show here, for the first time, that following systemic administration (i.e. intraperitoneally) CpG-C was abundantly taken up by TLR9-expressing cells across the brain, without affecting BBB integrity or infiltration of immune cells into the brain (Fig. 4, Supplementary Fig. 4), and dramatically reduced brain colonization by circulating tumor cells (Figs. 1,2,6,7, Supplementary Figs. 2,6). These findings pave the road for exploiting this compound in the clinic, as it could be easily administered systemically to serve as a prophylactic agent for patients with high risk of developing brain metastases.

An even more urgent clinical scenario where this treatment could prove life-saving is the perioperative period –days to weeks before and after tumor excision– which is now acknowledged as a critical therapeutic window for reducing post-operative metastatic disease (14,66). Indeed, various short perioperative interventions were reported to markedly impact short- and long-term cancer outcomes (14,67–69). As brain metastases are common in cancer patients and are associated with poor prognosis (1), reducing their post-operative occurrence is key in improving survival (4). Here, we show that in a spontaneous brain metastasis model of melanoma, a short perioperative treatment with CpG-C, spanning three days prior and following primary tumor excision, results in reduced brain tumor burden (Fig. 1g). Importantly, CpG-C was shown to have negligible toxicity in humans (19–21). While we did not directly test whether systemic CpG-C administration has any deleterious effects on neuronal activity, it has been shown by others that when administered directly into the brain (resulting in higher local concentrations) CpG-ODNs do not cause neurotoxicity in animals (27), nor result in significant or permanent neurological deficits in humans (19–21). Therefore, while traditional chemo and radiation therapies cannot be used during the perioperative period (due to their deleterious effects on tissue healing and immune competence), the use of CpG-C could be a promising prophylactic approach during this critical timeframe (14).

In pre-clinical trials, acute and chronic systemic CpG-ODNs (including CpG-C) were shown to reduce primary tumor growth and metastases in peripheral organs (10–12,70). Importantly, CpG-ODNs are evaluated as stand-alone anti-tumor agents as well as vaccine adjuvants in several clinical trials of different cancers, and systemic administration is considered well tolerated with negligible toxicity (71,72). Given the low toxicity of CpG-C and its wide-range anti-tumor effects, extended use beyond the perioperative period can also be considered. Additionally, TLR9 stimulation of microglia cells has also been shown to be beneficial in various neurological pathologies, including Alzheimer’s (26) and seizure-induced aberrant neurogenesis (28), although systemic treatment has not been studied for these conditions. As such, systemic CpG-C treatment could be considered as a therapeutic intervention for cancer and non-cancer-related pathologies.

It is well established that innate immune cells play a key role in preventing and eradicating metastases in the periphery (73–75). Indeed, we herein show that depletion of NK cells and monocytes results in elevated tumor-seeding in the lungs (Fig. 2a,c). However, in the brains of the same animals, we made the novel observation that NK and monocyte depletion has no effect, and that they do not mediate the beneficial effects of CpG-C (Fig. 2b,d). While mature NK cells are abundant in the capillaries of the lungs and liver (76), only limited numbers of immature NK cells are found in cerebral capillaries (77). Also, while patrolling monocytes (21) and pulmonary-resident macrophages are the first line of defense in the lungs (78), monocytes infiltrate the brain parenchyma only under pathological conditions in which the BBB is compromised (79,80), a condition that does not characterize the early stages of tumor cell infiltration (81). Notably, systemic CpG-C administration did not affect infiltration of T-cells (i.e. CD4^+^) or monocytes (i.e. CD68^+^) into the brain (Fig. 4d), or the number of GFP^+^ cells (i.e. monocytes/microglia) evident in the brain twenty-four hours following administration of tumor cells (Fig.6f). These differences between the periphery and the brain underscore the importance of studying brain-specific mechanisms that regulate the metastatic process, to allow tailoring of relevant therapies.

In the brain, microglia are the primary immune effector cells (53). Close interactions between macrophages/microglia cells and established metastases has been reported in human brain samples (82,83). In mice, it has been shown that heterogeneous microglia cells, activated and non-activated, accumulate proximal to invading tumor cells (55) and infiltrate established metastases generated by intracranial injection (84). However, the role of microglia in regulating brain tumor progression, especially during the initial steps of tumor colonization, remains unclear (83,85–88). Notably, established tumors can modulate activation of microglia, recruiting them to support tumor progression, whereas enabling microglia activation has an opposite effect (87–90). Our *in vitro* results indicate that both primary cultured and N9 microglia cells exert low tumoricidal activity (Fig. 5), in line with previous findings (91). Here, however, we clearly show that activation of microglia with CpG-C markedly increases this cytotoxic activity, mediated through direct physical contact with tumor cells and not in a paracrine fashion. While it has been argued that microglia cells promote initial steps of colonization of breast tumor cells *in vitro* and in acute slices (83), we found through *in vivo* two-photon imaging that microglia contact and phagocytize tumor cells immediately after their infiltration into the brain (Fig. 6a-c), and do so more abundantly following systemic administration of CpG-C (Fig. 6d,e). Accordingly, CpG-C increased mRNA expression of apoptosis-inducing and phagocytosis-related genes in microglia (Fig. 8), without affecting microglia density (Fig. 6f). Further, by blocking microglia activation (Fig. 7a,b), and by depleting them (Fig. 7c), we show that microglia mediate the beneficial *in vivo* anti-metastatic effects of CpG-C.

The metastatic process involves several steps, including arrest in the brain vasculature, infiltration through the BBB (mainly) and colonization of the brain parenchyma (92). Although CpG-C could have affected all of these steps in different magnitudes as endothelial cells and astrocytes also uptake the adjuvant (Fig. 3), we clearly show that the pool of metastatic cells infiltrating the brain was not altered (Fig. 6g) leading to the conclusion that, even if not directly measured, arrest and infiltration were not significantly affected by CpG-C. Support for this argument comes also from our findings that the permeability of key brain-immune interfaces was not altered (Fig. 4 and Supplementary Fig. 4). This conclusion does not overrule contribution of other mediators, which will become the topic of future research.

Overall, we demonstrate that shifting the balance from non-activated to activated microglia, as with the systemic CpG-C treatment presented herein, results in killing of invading tumor cells and prevents establishment of brain metastases. Such an approach could lay the foundation for a novel clinical perioperative therapy.

## Acknowledgement

This study was supported by a NIH/NCI R01 CA172138 (SBE) and the Leducq Foundation – Protect Stroke 15CVD02 (PB). AB is grateful to Dante family and Sagol School of Neuroscience of Tel Aviv University for the award of doctoral fellowship. NE wants to acknowledge support from the Melanoma Research Alliance (the Saban Family Foundation–MRA Team Science Award). The authors thank Reuven Stein and Uri Amit for commenting on early versions of this manuscript.

## Author Contributions

Conceptualization, A.B., L.M., S.B-E. and P.B.; Methodology, A.B., A.C., S.B-E. and P.B.; Software, P.B.; Investigation, A.B., M.G., A.L., D.K., A.C., M.A., and L.S.; Resources, A.G.; Writing – Original Draft, A.B., S.B-E. and P.B.; Writing – Review & Editing, A.B., M.G., M.A., N.E., L.M., S.B-E., and P.B.; Visualization, A.B. and P.B.; Supervision, N.E., D.A., L.M., S.B-E., P.B.

## Declaration of Interests

The authors declare no competing interests.

---

**Supplementary figure 1:**
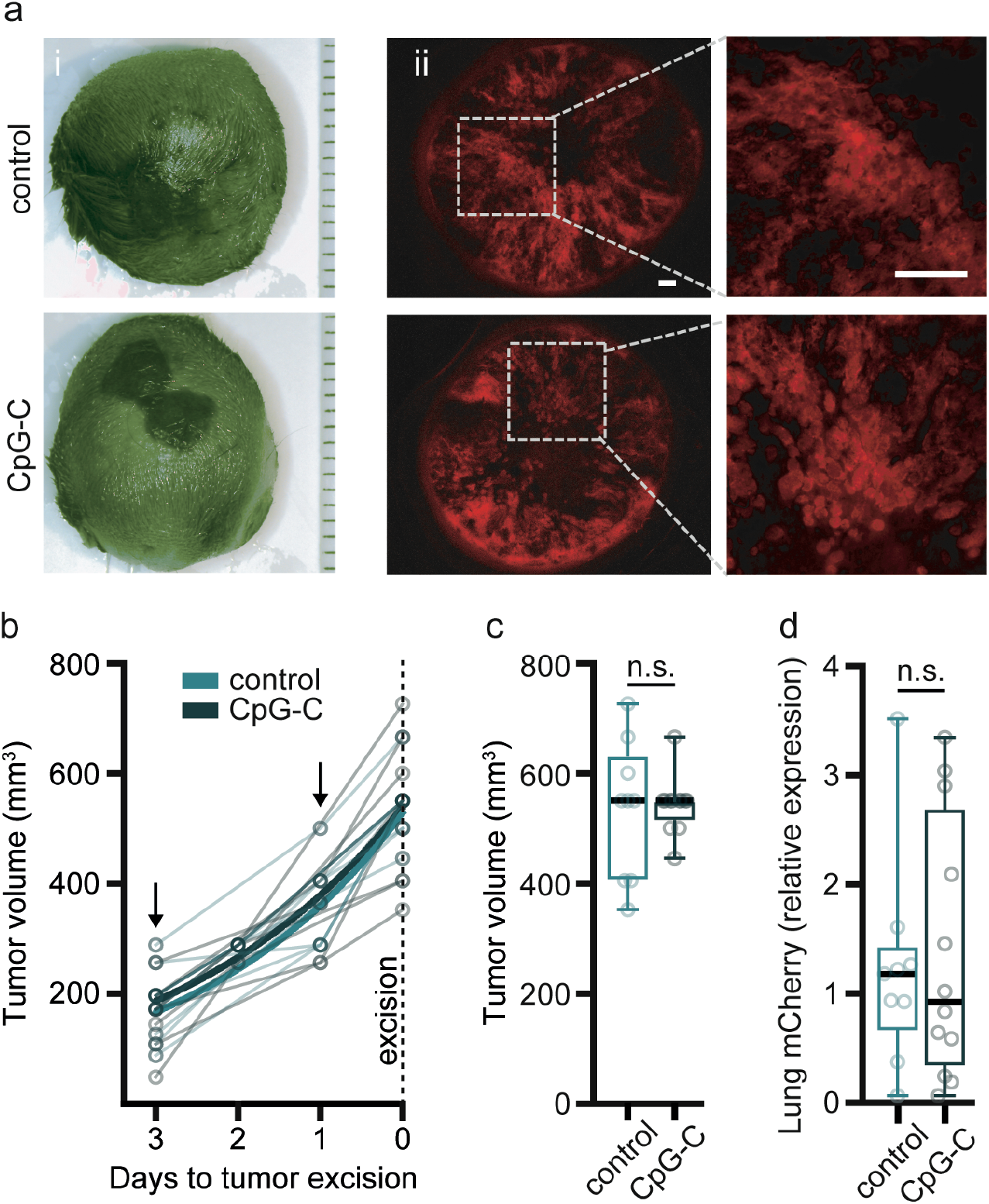
Perioperative CpG-C treatment did not affect primary melanoma tumor growth or spontaneous lung metastasis. (**a**) Representative images of melanoma Ret-mCherry primary tumor mass (left panels) and sections (right panels) from control and CpG-C treated animals. No differences in tumor appearance were evident. (**b-c**) CpG-C treatments (arrows) did not affect primary tumor growth dynamics (F_(2,60)_=0.5041, p=0.6066; for Y=Y0▯exp(k▯X) the 95% confidence intervals are: Y0=471.8 to 585.3, k=0.2890 to 0.4971, and Y0=509.0 to 571.1, k=0.3037 to 0.4089 for control and CpG-C, respectively; **b**). Tumors were excised from control and CpG-C treated animals at the same size (n=9 and n=12 for control and CpG-C, respectively; two-tailed Mann-Whitney U=52.50, p=0.9260; **c**). (**d**) CpG-C treatment during seven perioperative days did not affect micrometastases in the lung (measured by mCherry mRNA expression; n=9 and n=12 for control and CpG-C, respectively; two-tailed unpaired student t-test, t(19)=0.2756, p=0.7858). Data in (**b**) is presented as mean (±SEM) and boxplot whiskers represent min-max range (**c-d**).

**Supplementary figure 2:**
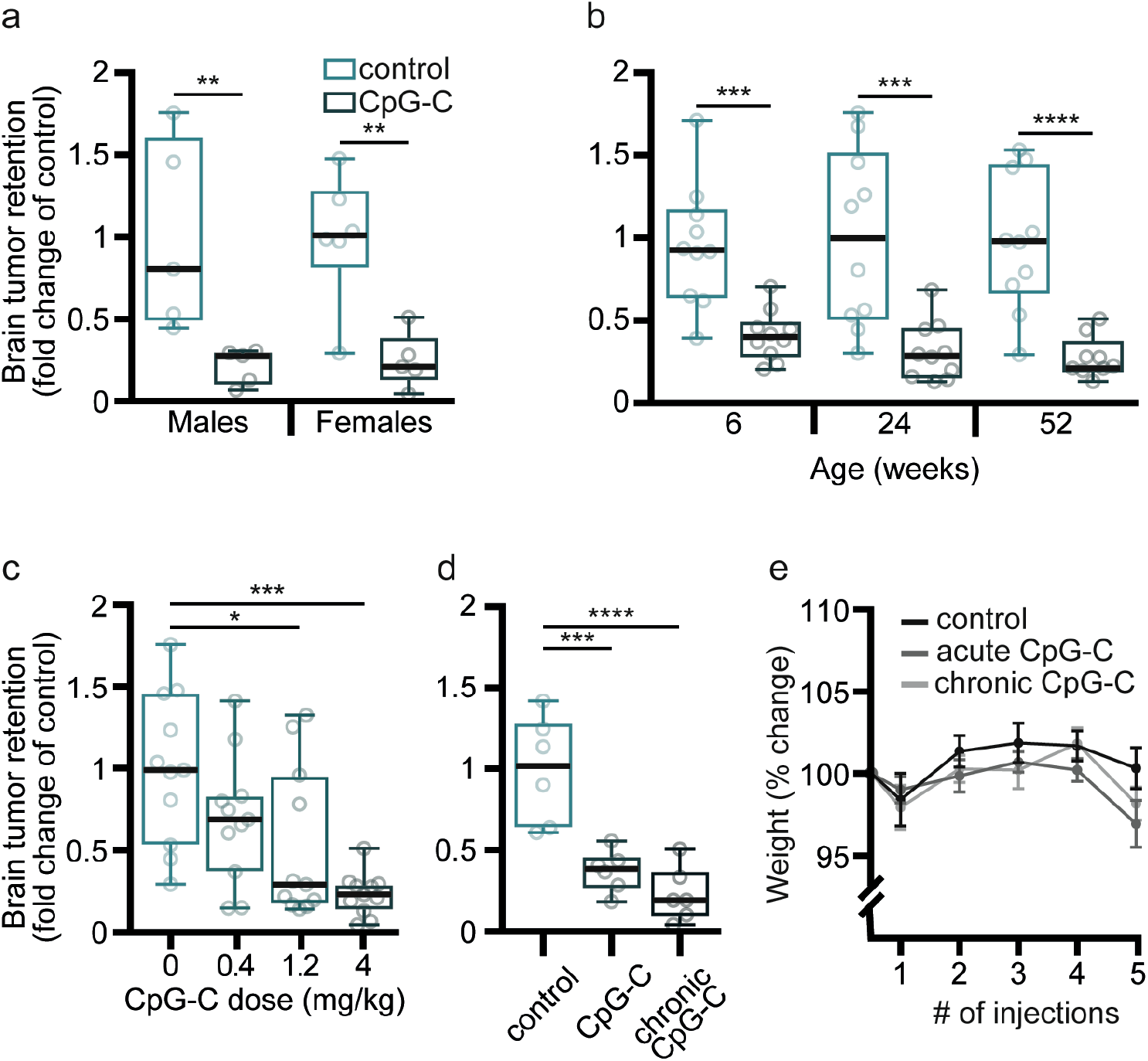
CpG-C is effective in reducing brain tumor retention in both sexes, across ages, in a dose-dependent manner, and both as an acute and as a chronic prophylactic treatment. (**a**) A systemic prophylactic injection CpG-C reduced brain tumor retention of D122 cells in both male (n=5, two-tailed Mann-Whitney U=0, p=0.0079) and female (n=5-6, two-tailed Mann-Whitney U=1, p=0.0087) mice to a similar degree. (**b**) CpG-C reduced brain tumor retention across ages - 6 weeks (n=10, two-tailed Mann-Whitney U=7, p=0.0005); 24 weeks (n=10, two-tailed Mann-Whitney U=8, p=0.0007); and 52 weeks (n=10, two-tailed Mann-Whitney U=2, p<0.0001). (**c**) CpG-C reduced brain tumor retention in a dose dependent manner (n=10-11, Kruskal-Wallis H=15.98, p=0.0011) reaching significance at 1.2mg/kg (p=0.0455), and with higher efficacy at 4mg/kg (p=0.0003). (**d**) An acute systemic injection of CpG-C one day before tumor cell injection (p=0.0298) was effective as chronic injections (every other day, starting ten days before tumor inoculation; p=0.0013) in reducing brain tumor retention (n=6, Kruskal-Wallis H=12.33, p=0.0001). (**e**) No weight loss was evident in animals receiving either acute or chronic systemic CpG-C treatment (n=6, two-tailed two-way ANOVA; F_(2,17)_=1.463, p=0.2593). Boxplot whiskers represent min-max range (**a-d**) and data in (**e**) is presented as mean (±SEM).

**Supplementary figure 3:**
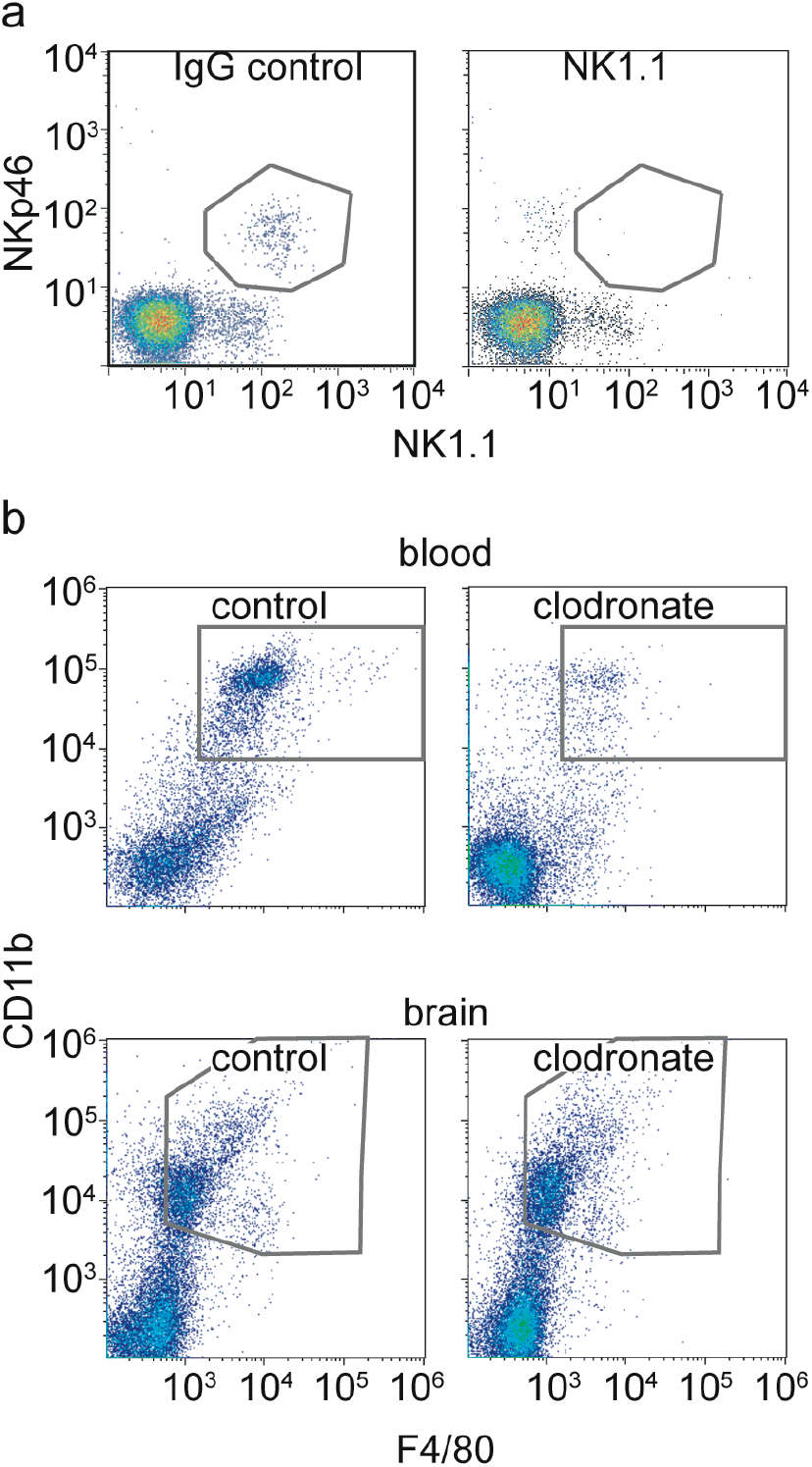
NK and monocyte depletion. (**a**) anti-NK1.1 injection resulted in >90% depletion of NK cells from the blood compared to IgG control. (**b**) Clodronate liposomes resulted in >85% depletion of monocytes from the blood (top panels), without affecting microglia viability (lower panels).

**Supplementary figure 4:**
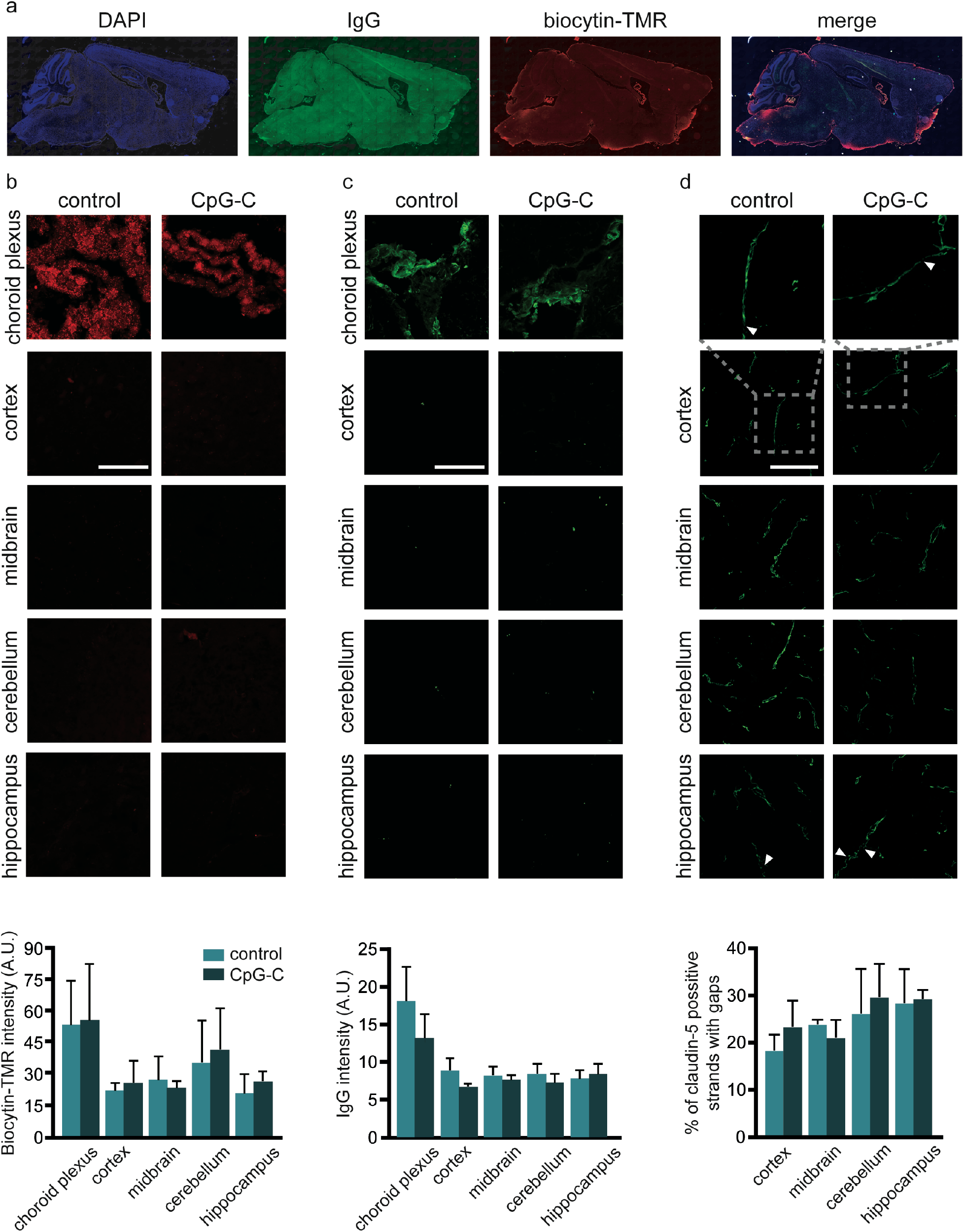
CpG-C does not affect BBB integrity. Mice (n=3) were treated with a single systemic (i.p.) injection of CpG-C (4mg/kg), and 24 hours later biocytin-TMR and IgG infiltration and claudin-5 continuity were measured in the cortex, cerebellum, midbrain, and hippocampus (five images for each anatomical region; see methods). (**a**) A tiled sagittal section of a CpG-C treated mouse. (**b-d**) CpG-C treatment did not affect blood vessels leakiness (F_(1,20)_=0.0828, p=0.7765 and F_(1,20)_=1.738, p=0.2023 for biocytin-TMR and IgG, respectively; **b-c**); nor claudin-5 continuity (F_(1,11)_=0.1272, p=0.7281; **d**) in any of the analyzed brain regions. Scale bar is 50μm. Data is presented as mean (±SEM).

**Supplementary figure 5:**
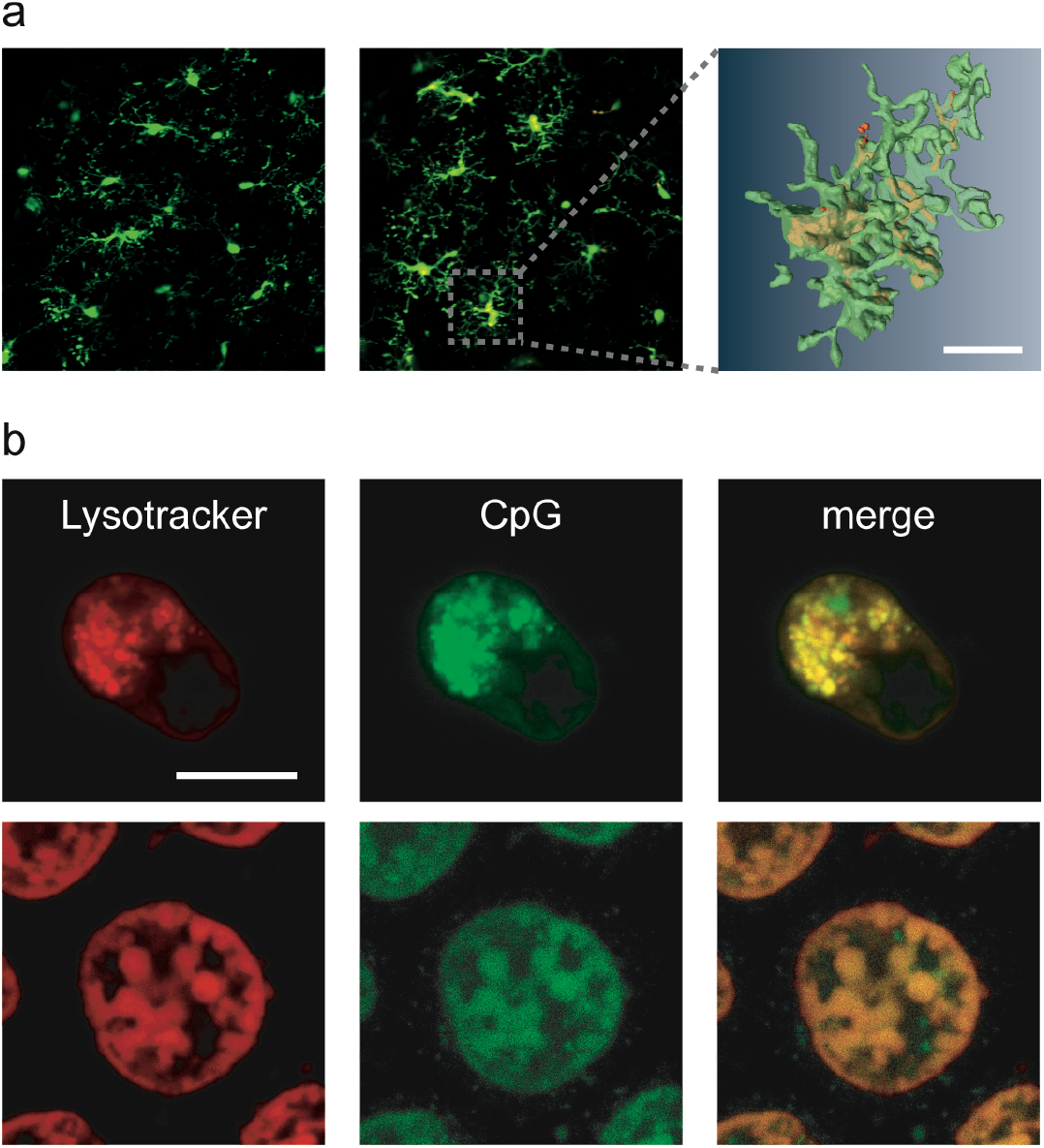
CpG-C is taken up into microglia lysosomes *in vitro* and *in vivo*. (**a**) TAMRA-labeled CpG-C injected systemically is taken up by microglia *in vivo* in CX3CR1^GFP/+^ mice (top left - before CpG-C injection; bottom left - after CpG-C injection; right panel - partial reconstruction; 15μm stacks, with 1μm z-steps). (**b**) N9 cells pretreated with TAMRA-labeled CpG-C for 24 hours (top panels) and microglia cells extracted from CX3CR1^GFP/+^ mice that were injected with TAMRA-labeled CpG-C 24 hours earlier (bottom panels) were co-stained with Lysotracker, demonstrating CpG-C was taken up into the lysosomes.

**Supplementary figure 6:**
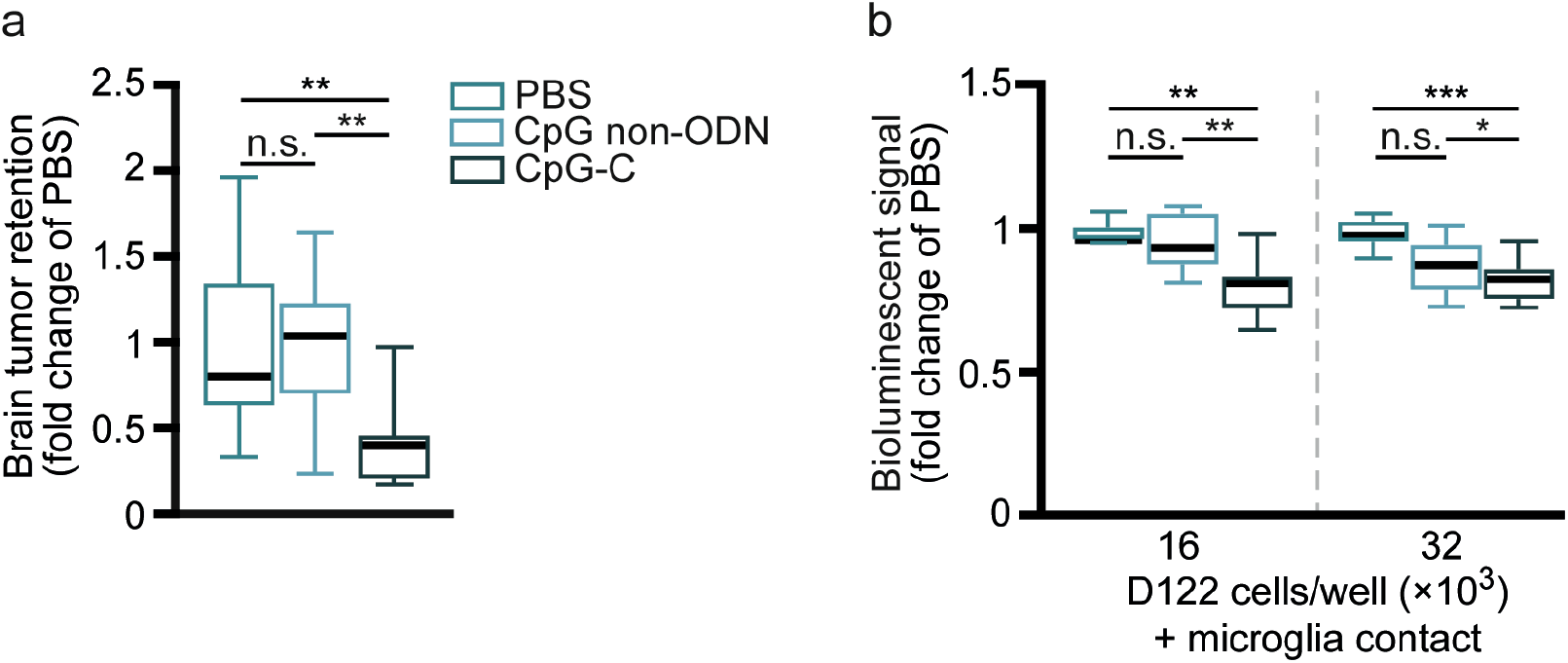
PBS and non-CpG ODN affect tumor cells viability similarly. (**a**) No differences in brain tumor retention were evident between PBS and non-CpG ODN treated animals (p=0.9974), while CpG-C significantly reduced brain tumor retention of D122 cells (F_(2,28)_=8.277, p=0.0040 and p=0.0048 compared to PBS and non-CpG ODN, respectively). (**b**) D122 cells were co-cultured in contact with N9 cells treated with PBS, non-CpG ODN, or CpG-C. No differences in tumor cells viability were evident between PBS and non-CpG ODN treated cultures (p=0.7745 and p=0.1420 for 16▯10^3^ and 32▯10^3^ D122 cells/well), while CpG-C significantly reduced tumor cells viability (for 16▯10^3^: F_(2,20)_=9.767, p=0.0017 and p=0.0062 compared to PBS and non-CpG ODN, respectively, and for 32▯10^3^: F_(2,19)_=12.15, p=0.0003 and p=0.0477 compared to PBS and non-CpG ODN, respectively). Boxplot whiskers represent min-max range.

**Supplementary figure 7:**
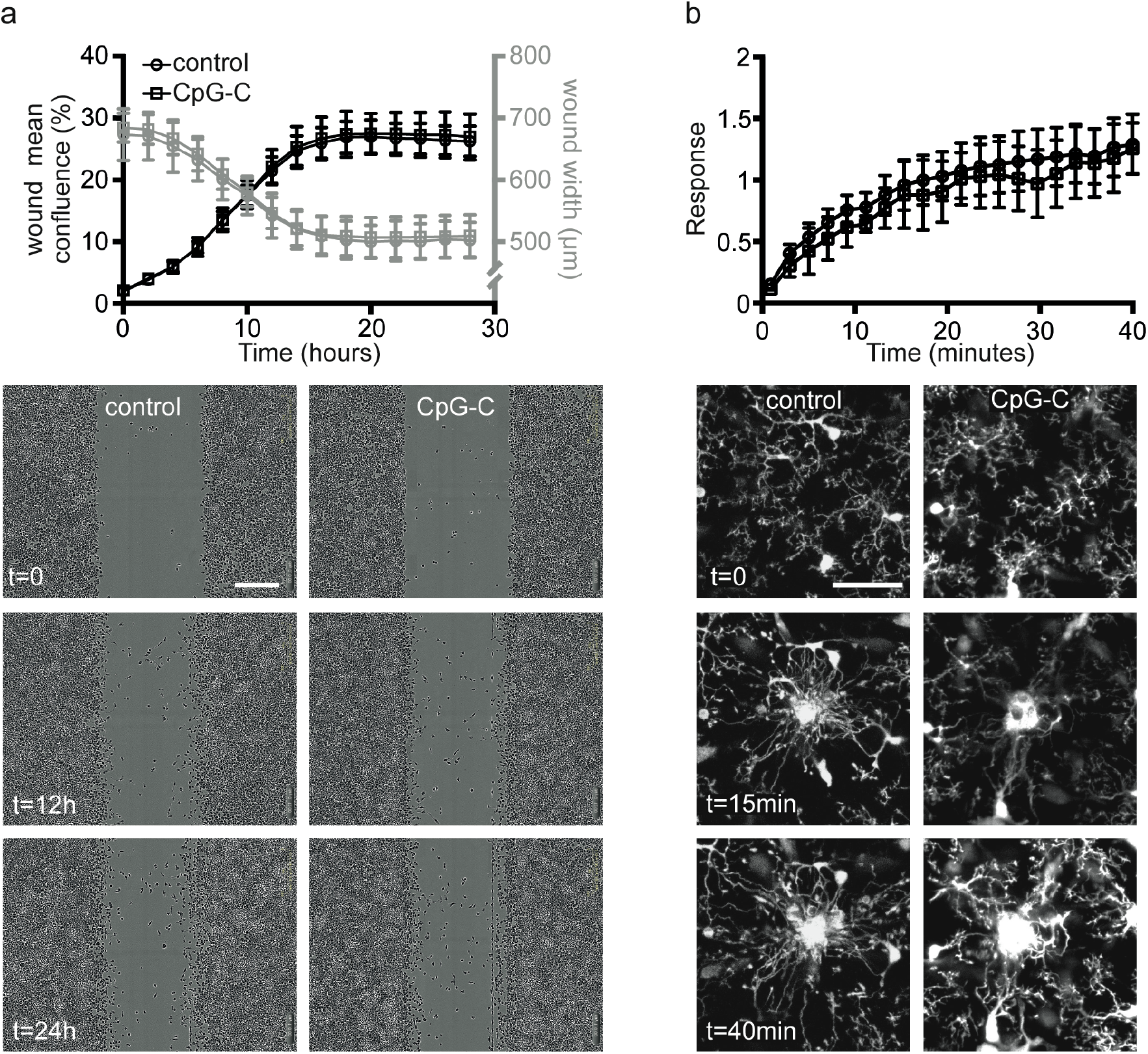
CpG-C does not affect microglia reaction to non-tumor related insults. (**a**) Microglial N9 cultures treated with 100nM/L CpG-C for 24 hours reacted similarly in the scratch migration assay compared to cultures treated with non-CpG ODN, indicated by wound confluence (F_(1,16)_=0.1845, p=0.6732) and wound width (F_(1,16)_=0.2801, p=0.6039). Scale bars is 300μm. (**b**) Microglia reacted similarly to a photodamage induced *in vivo* by a high-power laser (780 nm; 150mW at the sample; ~1μm size) in CpG-C treated and control CX3CR1^GFP/+^ mice (F_(1,8)_=0.1111, p=0.7474). Scale bars is 50μm. Data is presented as mean (±SEM).

**Supplementary figure 8:**
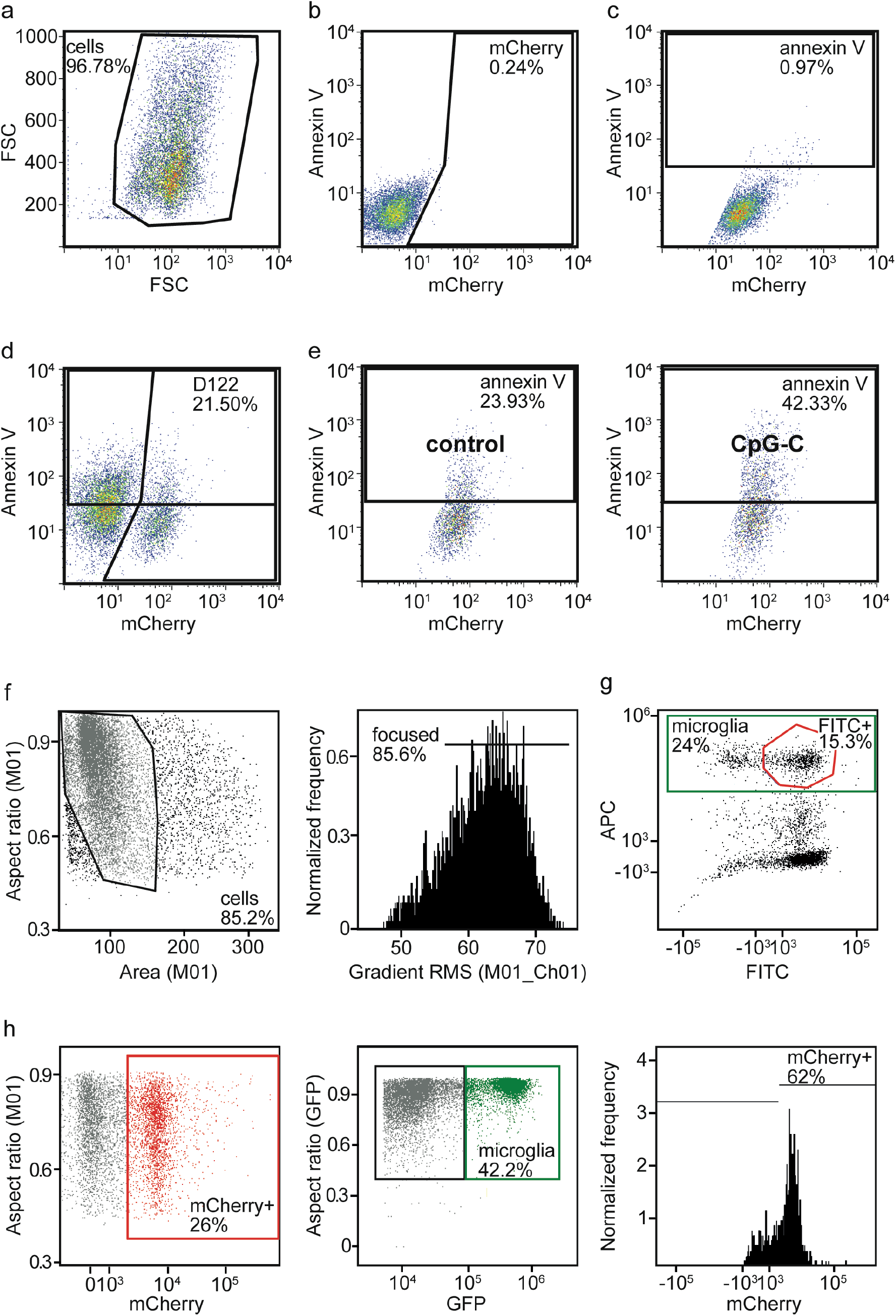
FACS analyses gating strategies. (**a**) Annexin V *in vitro* experiments (Fig. 4h) were analyzed by selecting single cells from a plot of SSC against FSC. (**b-d**) The gates for mCherry positive cells (D122; **b**) and annexin V positive cells (**c**) were selected based on samples negative for these staining and validated using positive control samples containing N9 and mCherry-labeled D122 cells (**d**). To induce annexin V staining (indicating apoptosis) in positive control samples, half of the cells were placed in 90 degrees Celsius for 2 min and then immediately on ice for 2 minutes and mixed together. (**e**) Examples for CpG non-ODN (control; left panel) and CpG-C treated wells (right panel). (**f**) ImageStream image data files were analyzed by selecting single cells from a plot of object area against object aspect ratio (width/length; left panel) and then focused cells using the Gradient RMS feature (right panel). (**g**) For CpG-C-uptake experiments (Fig. 3c), cells that have taken up FITC-labeled CpG-C were identified on a scatter plot of FITC against the appropriate fluorophore (e.g. APC for microglia cells). (**h**) For experiments in (Fig. 5e-g,i) mCherry positive cells (left panel) and microglia cells (from CX3CR1^GFP/+^ mice; middle panel) were identified on a scatter plot of intensity of the relevant fluorophore against object aspect ratio. mCherry positive microglia cells were identified inside the microglia sub-population in a histogram of the of mCherry intensity (right panel). For quantification of (**g**) and (**h**; right panel) we used the internalization wizard.

**Supplementary movie:**

Microglia (white) treated with CpG-C phagocytize invading tumor cells (red) *in vivo* as early as few hours after tumor cell inoculation. Orange arrows mark phagocytosis events at day 0 (left) and their corresponding events at day 1 (right). Field of view for each day is 200μm.

